# Density-dependent resistance protects *Legionella pneumophila* from its own antimicrobial metabolite, HGA

**DOI:** 10.1101/383018

**Authors:** Tera C. Levin, Brian P. Goldspiel, Harmit S. Malik

**Affiliations:** Division of Basic Sciences, Fred Hutchinson Cancer Research Center, Seattle WA; Howard Hughes Medical Institute, Fred Hutchinson Cancer Research Center, Seattle WA

## Abstract

To persist in the extracellular state, the bacterial pathogen *Legionella pneumophila* must withstand competition from neighboring bacteria. Here, we find that *L. pneumophila* can antagonize the growth of neighboring *Legionella* species using a secreted inhibitor: HGA (homogentisic acid), the unstable, redox-active precursor molecule to *L. pneumophila*’s brown-black pigment. Unexpectedly, we find that *L. pneumophila* can itself be inhibited by HGA secreted from neighboring, isogenic strains. Our genetic approaches further identify *lpg1681* as a gene that modulates *L. pneumophila* susceptibility to HGA. We find that *L. pneumophila* sensitivity to HGA is density-dependent and cell intrinsic. This resistance is not mediated by the stringent response nor the previously described *Legionella* quorum-sensing pathway. Instead, we find that *L. pneumophila* cells secrete HGA only when they are conditionally HGA-resistant, which allows these bacteria to produce a potentially self-toxic molecule while restricting the opportunity for self-harm. We speculate that established *Legionella* communities may deploy molecules such as HGA as an unusual public good that can protect against invasion by low-density competitors.

## Introduction

Inter-bacterial conflict is ubiquitous in nature, particularly in the dense and highly competitive microenvironments of biofilms (Davey and O’toole 2000; Foster and Bell 2012; Ghigo and Rendueles 2015). In these settings, bacteria must battle for space and nutrients while evading antagonism by neighboring cells. One strategy for managing these environments is for bacteria to cooperate with their kin cells, sharing secreted molecules as public goods (Nadell, Drescher, and Foster 2016; Abisado et al. 2018). However, these public goods are vulnerable to exploitation by other species or by ‘cheater’ bacterial strains that benefit from public goods but do not contribute to their production. For this reason, many bacteria participate in both cooperative and antagonistic behaviors to survive in multispecies biofilms. Bacterial antagonistic factors can range from small molecules to large proteins, delivered directly or by diffusion, and can either act on a broad spectrum of bacterial taxa or narrowly target only a few species. Although narrowly targeted mechanisms may seem to be of less utility than those that enable antagonism against diverse bacterial competitors, targeted strategies can be critical for bacterial success because they tend to mediate competition between closely-related organisms that are most likely to overlap in their requirements for restricted nutrients and niches (Hibbing et al. 2010).

The bacterium *Legionella pneumophila* (*Lp*) naturally inhabits nutrient-poor aquatic environments where it undergoes a bi-phasic lifestyle, alternating between replication in host eukaryotes and residence in multi-species biofilms (Lau and Ashbolt 2009; Declerck et al. 2007; Declerck 2010; Taylor, Ross, and Bentham 2013). If *Lp* undergoes this lifecycle within man-made structures such as cooling towers or showers, the bacterium can become aerosolized and cause outbreaks of a severe, pneumonia-like disease in humans, called Legionnaires’ disease (Fraser et al. 1977; McDade et al. 1977; Fields, Benson, and Besser 2002). Because of the serious consequences of *Lp* colonization, the persistence and growth of *Legionella* in aquatic environments has been the subject of numerous studies. These studies have examined replication within protozoan hosts (Rowbotham 1980; Isberg, O’Connor, and Heidtman 2008; Lau and Ashbolt 2009; Hoffmann, Harrison, and Hilbi 2014), survival in water under nutrient stress (Li et al. 2015; Mendis, McBride, and Faucher 2015), and sensitivity to biocides (Kim et al. 2002; Lin, Stout, and Yu 2011). Here, we focus on interbacterial competition as an underappreciated survival challenge for *Lp*.

*Legionella spp.* are not known to produce any antibiotics, bacteriocins, or other antibacterial toxins. Bioinformatic surveys of *Legionella* genomes have revealed a number of polyketide synthases and other loci that likely produce bioactive metabolites (Johnston et al. 2016; Tobias et al. 2016), but these have not been shown to exhibit any antimicrobial functions. Nevertheless, there are some hints that interbacterial competition is relevant for *Lp* success within biofilms. For example, one study of artificial two-species biofilms found that viable *Lp* were able to persist for over two weeks in the presence of several bacterial species (e.g. *Pseudomonas fluorescens, Klebsiella pneumoniae*) but not others (e.g. *Pseudomonas aeruginosa*) (Stewart, Muthye, and Cianciotto 2012). Additionally, *Lp* bacteria are often co-resident with other *Legionella spp.* in man-made structures, with some studies showing that *Lp* proliferation is correlated with a decrease in other *Legionella spp.* populations (Wery et al. 2008; Pereira et al. 2017; Declerck et al. 2007). These studies suggest that *Lp* bacteria may compete with other *Legionella spp.* for similar biological niches.

The most direct evidence for interbacterial competition comes from Stewart et al., 2011, who found that *Lp* could antagonize the growth of neighboring *Legionella spp.* on the same plate (Stewart, Burnside, and Cianciotto 2011). The molecules mediating this competition have not been identified, although previous work suggested a role for *Lp*’s secreted surfactant, a thin liquid film that facilitates the spread of *Lp* across agar plates (Stewart, Rossier, and Cianciotto 2009; Stewart, Burnside, and Cianciotto 2011). Still, it remained unknown if surfactant played a direct or indirect role in inter-*Legionella* inhibition.

Here, we use unbiased genetic approaches to find that homogentisic acid (HGA) produced by *Lp* inhibits the growth of neighboring *Legionella spp*. We find that HGA production co-occurs with surfactant production, but that these are independent, separable phenomena. The redox state of HGA appears to be critical for its activity, as oxidized HGA-melanin pigment is inactive. Unexpectedly, we find that *Lp* itself is susceptible to HGA inhibition. We also identify one gene– *lpg1681*– that enhances *Lp* susceptibility to HGA. However, we find that *Lp* cells can be resistant to HGA at high-density, which is also when they secrete large amounts of HGA. This high-density resistance is cell intrinsic and appears to be independent of growth phase, the stringent response or the previously described quorum-sensing pathway in *Legionella*. Based on these findings, we propose that HGA has the potential to play an important role in structuring *Legionella* communities.

## Results

### *L. pneumophila* inhibits *L. micdadei* via an unknown, secreted inhibitor

Inspired by previous reports (Stewart, Burnside, and Cianciotto 2011), we investigated how *Legionella pneumophila* (*Lp*) engages in inter-*Legionella* competition. We found that *Lp* inhibited the growth of neighboring *Legionella micdadei* (*Lm*) plated 1 cm away on solid media, suggesting that it produces a secreted inhibitor (Figure 1A). This inhibition was most robust when we plated the *Lp* strain on low-cysteine media 3-4 days prior to plating *Lm*, allowing time for the inhibitory molecule to be produced and spread across the plate. To quantify this inhibition, we recovered *Lm* grown at different distances from *Lp*. After 48h incubation, we found a 10,000-fold difference in growth between *Lm* antagonized by *Lp* versus *Lm* plated outside of the zone of inhibition (Figure 1B).

**Figure 1:**
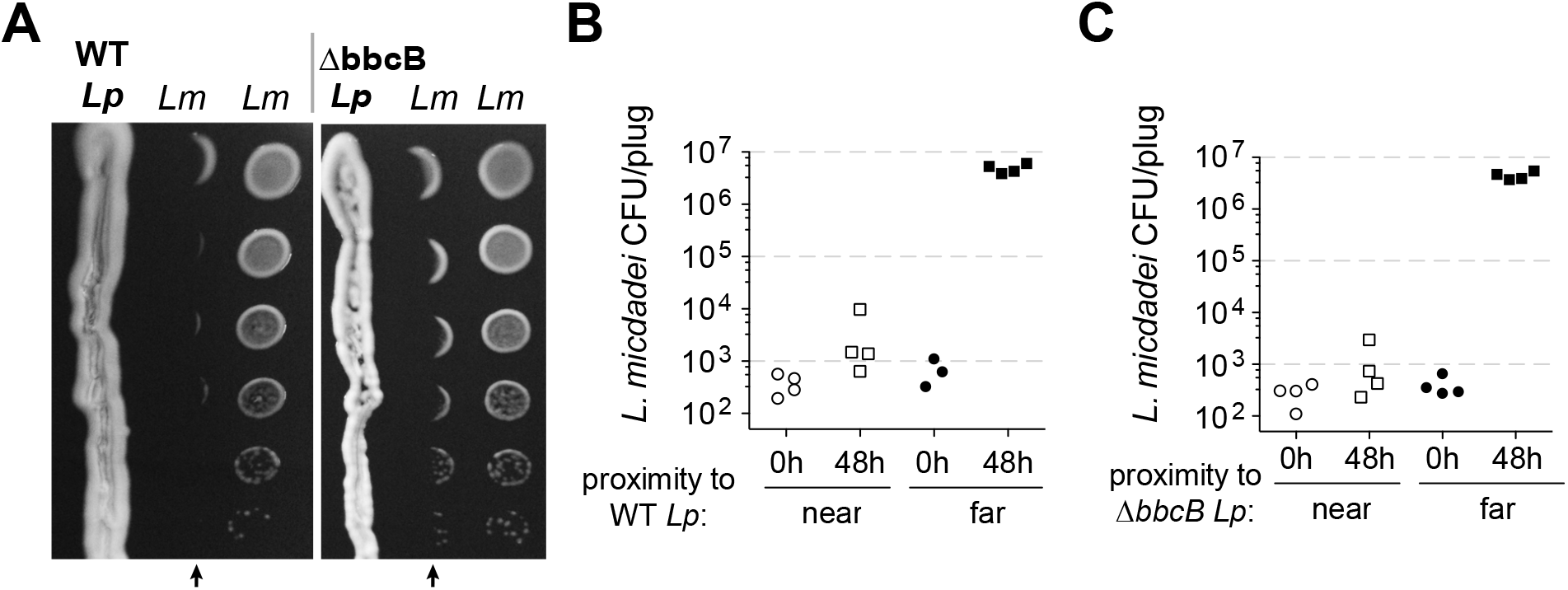
*L. pneumophila (Lp)* produces a secreted inhibitor independent of surfactant. A) When pre-incubated on BCYE charcoal agar plates, *Lp* produces a zone of inhibition, impacting the growth of nearby *L. micdadei (Lm)*. Arrows mark the edge of inhibition fronts. Droplets of *Lm* at different dilutions were added to the plate 3 days after streaking *Lp*. Outside of the zone of inhibition, *Lm* grows in a circle where spotted on the plate, while inhibition either prevents growth completely or results in a crescent of growth away from *Lp*. WT *Lp* (left panel) generates a similar zone of inhibition as a surfactant-null mutant, Δ*bbcB* (right panel). B) Quantification of *Lm* growth within (”near”) or outside of (”far”) the wild type *Lp* zone of inhibition. C) Quantification of *Lm* growth within or outside of the Δ*bbcB Lp* zone of inhibition. In B and C bacteria were sampled and removed from the plate in a “plug” of fixed area before plating for viable CFUs.

Previous studies (Stewart, Burnside, and Cianciotto 2011) had proposed that inter-*Legionella* inhibition could be caused by *Lp*’s secreted surfactant, which is produced by *Lp* but not *Lm* (Stewart, Rossier, and Cianciotto 2009). We tested this hypothesis by deleting a surfactant biosynthesis gene, *bbcB*, from the *Lp* genome (Stewart, Burnside, and Cianciotto 2011). The resulting Δ*bbcB* strain did not produce surfactant (Supplemental figure 1B), yet it still inhibited adjacent *Lm* (Figure 1A, Supplemental Figure 1C). When quantified, we observed nearly identical inhibition from both wild type *Lp* and Δ*bbcB Lp* (Figure 1C) indicating that the surfactant did not enhance inhibition levels. Furthermore, we observed that the zone of inhibition surrounding wildtype *Lp* did not always co-occur with the spread of the surfactant front (Supplemental figure 1A). From these results, we conclude that *L. pneumophila* can cause strong growth inhibition of neighboring *Legionella* using an unknown molecule that is distinct from surfactant.

### Transposon screen pinpoints HGA-melanin pathway in inter-*Legionella* inhibition

To determine which molecule(s) might be responsible for inter-*Legionella* inhibition, we performed an unbiased genetic screen in *Lp*. We generated *Lp* mutants using a drug-marked Mariner transposon that randomly and efficiently integrates into the *Legionella* genome (O’Connor et al. 2011). To identify mutants that were defective in producing the inhibitor, we transferred each mutant onto a lawn of *L. micdadei* on low-cysteine plates and examined the resulting zone of inhibition surrounding each *Lp* mutant (Figure 2A). After screening 2870 clones, we isolated 19 mutants that produced a smaller zone of inhibition than wild type *Lp*, as well as 5 mutants that showed a complete loss of inhibition (Figure 2B, Supplemental Table 1). We refer to these as “small zone” and “no zone” mutants, respectively. Among the “small zone” mutants, some had defects in surfactant spreading on plates, while others enhanced surfactant spread (Supplemental figure 2A), further distinguishing inter-bacterial inhibition from surfactant secretion.

**Figure 2:**
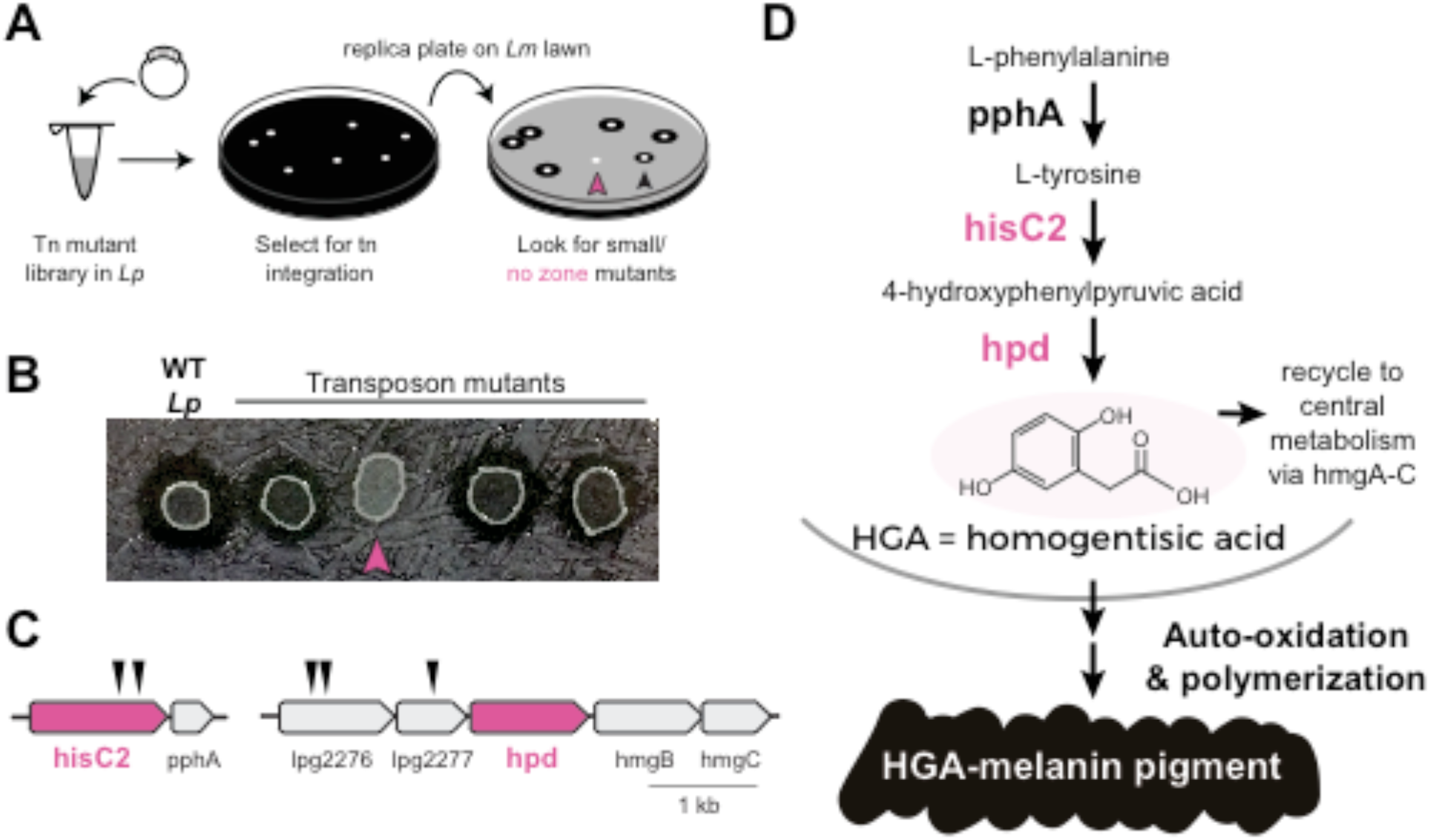
Transposon mutagenesis screen implicates HGA-melanin pathway in production of inhibitor. **A)** Screen for mutant *L pneumophila* (*Lp)* that do not inhibit *L. micdadei (Lm).* Following electroporation of a Mariner-transposon-containing plasmid, *Lp* mutants were selected for transposon integration. Colonies were patched or replica plated onto a lawn of *Lm.* Mutants of interest generated a zone of mhibrtran that was reduced (black arrowhead) or absent (pink arrowhead) compared to WT *Lp.* **B)** Selected transposon mutants produce abnormal zones of inhibition when grown on a lawn of *Lm.* Pink arrowhead indicates the ft1sC2::Tn “no zone” mutant. **C)** Transposon insertion sites (triangles) identified in the five recovered “no zone” mutants. **D)** HGA-melanin synthesis pathway. HGA is exported from the cell where it auto-oxidizes and polymerizes to form HGA-melanin. Genes indicated in pink were validated by complementation to have essential roles in *Lm* inhibition.

We focused on the “no zone” mutants, as these had the strongest defects in inhibition. These 5 mutants carried transposon insertions in two separate operons (Figure 2C). The first operon had two insertions in the *hisC2* gene (*lpg1998)*, which breaks down tyrosine as part of the HGA-melanin metabolic pathway (Figure 2D). Its downstream gene, *pphA*, converts phenylalanine to tyrosine in the same pathway. To validate the role of *hisC2* in inhibition, we overexpressed this gene in the *hisC2* transposon mutant background and found that *hisC2* alone was sufficient to complement the mutant phenotype (Supplemental Figure 2B). Having confirmed the role of *hisC2*, we turned to the second operon, where we had recovered transposon insertions in two uncharacterized genes, *lpg2276* and *lpg2277* (Figure 2C). These two genes lie immediately upstream of *hpd* (*lpg2278*), which is known to act with *hisC2* in the HGA-melanin pathway (Steinert et al. 2001; Gu et al. 1998) (Figure 2D). Because transposon insertions at the beginning of an operon can disrupt the expression of downstream genes via polar effects, we hypothesized that the insertions we recovered in *lpg2276* and *lpg2277* altered inter-*Legionella* inhibition via disruption of *hpd* expression. Indeed, we were able to complement insertions in both genes, which had yielded ‘no zone’ mutants, by overexpressing *hpd*, despite the fact that *hpd* overexpression caused a growth defect (Supplemental Figure 2B). In conclusion, all five “no zone” isolates had mutations that disrupted the same metabolic pathway involved in the production of HGA-melanin. Consistent with these findings, we observed defects in HGA-melanin pigmentation in all of the “no zone” mutants as well as some of the “small zone” mutants (Supplemental Figure 2E).

The HGA-melanin pathway is found in diverse eukaryotes and bacteria (Nosanchuk and Casadevall 2003; Liu and Nizet 2009) including *L. pneumophila* (Supplemental figure 2D). This pathway produces homogentisic acid (HGA) from the catabolism of phenylalanine or tyrosine (Fang, Yu, and Vickers 1989; Steinert et al. 2001) (Figure 2D). HGA can either be further metabolized and recycled within the cell via HmgA-C, or it can be secreted outside of the cell, where it auto-oxidizes and polymerizes to form a black-brown pigment called HGA-melanin, or pyomelanin (Kotob et al. 1995) (Figure 2D). To test whether intracellular metabolites downstream of HGA are necessary for inhibition, we deleted *hmgA*, the first gene in the pathway that recycles HGA back into central metabolism. We found that the Δ*hmgA* strain produced a zone of inhibition that was similar or slightly larger than wild type (Supplemental Figure 2C). We therefore inferred that synthesis of secreted HGA and/or HGA-melanin, but not its recycling and intracellular processing, is required for *Lp* inhibition of *Lm*.

### HGA inhibits the growth of *Legionella micdadei*, but HGA-melanin does not

To our knowledge, the HGA-melanin pathway has not previously been implicated in inter-bacterial competition. To the contrary, prior work has emphasized the beneficial (rather than detrimental) effects of HGA-melanin on *Legionella* growth, by providing improved iron scavenging (Chatfield and Cianciotto 2007) and protection from light (Steinert et al. 1995). We therefore asked whether the active inhibitor produced by the pathway was HGA-melanin, or alternatively a precursor molecule (Figure 3A). We tested the potential inhibitory activity of HGA-melanin pigment from *Lp* conditioned media in multiple experiments; however, we never observed any inhibition of *Lm*. We wished to rule out the possibility that the pigment secreted into rich media was too dilute to be active, or that other nutrients in the media might interfere with inhibition. We, therefore, isolated a crude extract of HGA-melanin from *Lp* conditioned media via acid precipitation (as in (Chatfield and Cianciotto 2007)), washed and concentrated the pigment approximately 10-fold and repeated the assay; the concentrated pigment also showed no inhibitory activity (Figure 3B, 3E).

**Figure 3:**
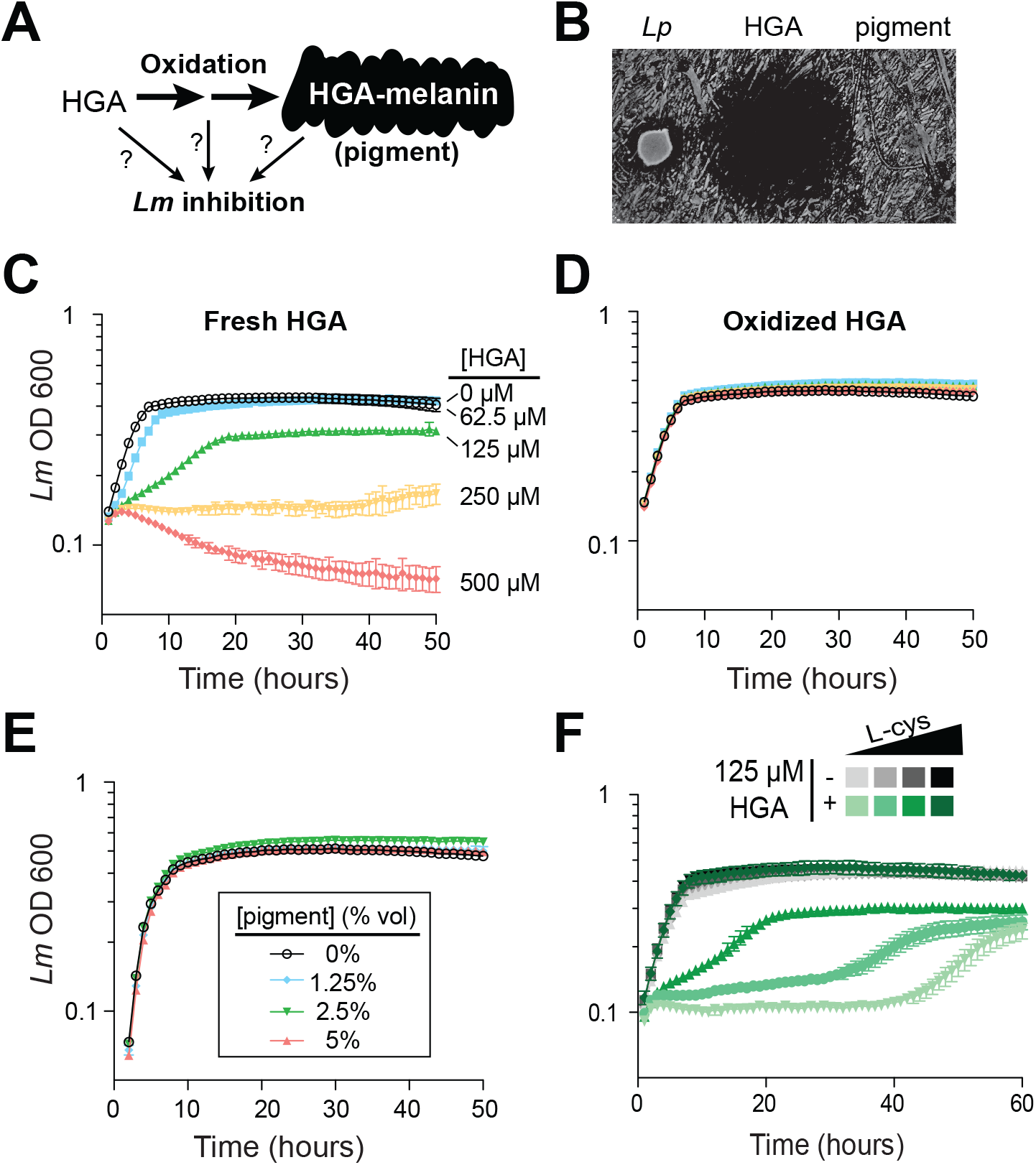
Synthetic HGA inhibits *Legionella micdadei* growth, depending on its oxidation state. **A)** We tested whether inter-bacterial inhibition is caused by HGA, HGA-melanin, or an oxidative intermediate. **B)** Zones of inhibition on a lawn of *Lm* generated from live *Lp* bacteria, synthetic 50mM HGA, or concentrated pigment extract. HGA prevents *Lm* growth in a large region (central dark circle) but pigment does not. **C)** Growth inhibition of *Lm* from increasing concentrations of synthetic HGA in rich AYE media. **D)** Pre-oxidation of synthetic HGA in AYE media for 24h eliminates its inhibitory activity, resulting in normal *Lm* growth. Concentrations of HGA colored as in panel C. **E**) Addition of oxidized HGA-melanin pigment from *L. pneumophila* has little impact on *Lm* growth in AYE liquid media. **F**) In the absence of HGA (gray symbols), titration of L-cysteine (L-cys) from 25%-200% of standard AYE media has little impact on *Lm* growth. However, HGA activity is enhanced in low-cysteine media and decreased in high-cysteine media (green symbols). All error bars show standard deviations among 3-4 replicates.

The first metabolite secreted by the HGA-melanin pathway is HGA. We tested whether HGA could behave as an inhibitor even though HGA-melanin could not. Indeed, we found that synthetic HGA robustly inhibited *Lm* growth, both when spotted onto a lawn of *Lm* and when titrated into AYE rich media (Figure 3B, 3C), We found that inhibition of *Lm* by HGA is relatively specific at the molecular level; neither 2- hydroxyphenylacetic acid nor 3-hydroxyphenylacetic acid, two HGA-related molecules that differ from HGA by only a single -OH group, were able to inhibit *Lm* growth at any concentration tested (Supplemental figure 3A).

Because HGA, but not HGA-melanin can inhibit *Lm* growth (Figure 3B, 3E), we inferred that the oxidative state of HGA might be important to its inhibitory activity. HGA is a reactive molecule, which auto-oxidizes (Eslami, Namazian, and Zare 2013) and polymerizes to form HGA-melanin through a series of non-enzymatic steps that are not genetically encoded (Steinert et al. 2001) and are therefore undetectable by our genetic screen. Given its auto-oxidative potential, we next tested whether HGA might cause growth inhibition by oxidizing other nearby molecules, either in the media or on bacterial cells. We allowed synthetic HGA to oxidize completely for 24h in AYE media before adding *Lm* (Supplemental Figure 4D). We found that pre-oxidation completely abolished synthetic HGA activity, even at very high HGA concentrations (Figure 3D, compare to 3C). This experiment also ruled out the possibility that HGA acts by causing nutrient depletion or other modifications of the media, since media pre-incubated with HGA for 24h is still able to support normal *Lm* growth. Instead, we infer that *Lm* inhibition results from direct interactions between bacterial cells and either HGA itself or unstable, reactive intermediates produced during HGA oxidation.

Small, reactive, quinone-like molecules similar to HGA are known to react with oxygen to produce H_2_O_2_, which is broadly toxic to bacteria (Hassan and Fridovich 1980). In such cases, extracellular catalase has been shown to protect bacteria against the toxic effects of H_2_O_2_ (Hassan and Fridovich 1980; Imlay 2013). To test if HGA toxicity occurs via a similar mechanism as H_2_O_2_, we asked if extracellular catalase was sufficient to protect *Lm* from HGA-mediated toxicity. Even at very high catalase concentrations, we found that catalase provided no protection from HGA (Supplemental Figure 3B), ruling out the production of extracellular H_2_O_2_ as a potential mechanism of action for HGA-mediated inhibition. We also considered the possibility that HGA as a weak acid could inhibit *Lm* indirectly by altering the local pH, but we observed that adding HGA at 1mM into AYE media or PBS caused little to no change in pH.

Given that the redox state of HGA is critical for inhibition, we reasoned that it should be possible to modulate HGA activity by altering the redox state of the media using reducing agents. We accomplished this by titrating L-cysteine from 25% to 200% of the levels in standard AYE media. In the absence of HGA, these altered cysteine concentrations had little impact on *Lm* growth (Figure 3F). However, lower cysteine concentrations greatly sensitized *Lm* to HGA, while excess cysteine was completely protective (Figure 3F). These findings may help partially explain why HGA’s inhibitory activity on *Legionella* has not been previously detected, as *Legionella* species are typically studied in cysteine-rich media. We found that synthetic HGA is readily able to react with cysteine in vitro (Supplemental Figure 3D) presumably impacting the oxidation state of HGA. Moreover, incubation of HGA with two other reducing agents, DTT (dithiothreitol) or reduced glutathione, similarly quenched HGA’s inhibitory activity (Supplemental Figure 3E). From these experiments, we conclude that HGA is less potent in rich media because it reacts with excess cysteine (or other bystander molecules) before it can interact with *Lm*. In sum, these results implicate the reactive activity of HGA and/or its transient, oxidative intermediates in inter-*Legionella* inhibition.

In these experiments, synthetic HGA was a robust inhibitor of *Lm.* However, this inhibition required relatively high concentrations of HGA (>50µM). We next quantified the amount of HGA secreted by *Lp* to determine if these levels were biologically relevant. Compared to other *Legionella spp.*, *Lp* produces much more pigment (Supplemental Figure 2D), suggesting that it secretes considerably more HGA. To estimate the quantity of secreted HGA, we created a standard curve of synthetic HGA in AYE rich media. We allowed the HGA added to completely oxidize to HGA-melanin, which can be measured by OD 400 readings. In this way, we can use the pigment levels after oxidation as a reliable measure of total HGA that was produced by a given time-point (Supplemental Figure 4D). Using this calibration, we estimated that wild type *Lp* had secreted the equivalent of 1.7mM HGA after 48h of culture, whereas the hyperpigmented Δ*hmgA* strain secreted about 2.6mM HGA. Thus, the levels of HGA produced by *Lp* are considerably higher than the inhibitory concentrations of synthetic HGA used in our assays (50-500 µM), at least under lab conditions. In contrast, we did not detect any pigment from the non-inhibitory *hisC2*::Tn strain (Supplemental figure 4E). From these experiments, we conclude that HGA is an abundant, secreted, redox-active metabolite of *Lp*, which can accumulate in concentrations that are relevant for inter-*Legionella* inhibition.

### *L. pneumophila* can be susceptible to its own inhibitor

Our results so far indicated that HGA can be a potent, redox-active inhibitor of *Lm*, which is volatile and capable of reacting with many types of thiol-containing molecules. If *Lp* uses HGA to compete with neighboring *Legionella spp.*, we anticipated that that *Lp* would have evolved some form of resistance to its own secreted inhibitor. Therefore, we next tested *Lp* susceptibility to inhibition in low-cysteine conditions, as we had previously done for *Lm*. Surprisingly, we found that *Lp* was quite sensitive to inhibition by neighboring *Lp* that was already growing on the plate (Figure 4A). Indeed, *Lp* susceptibility closely mirrored the susceptibility of *Lm* to inhibition (compare to Figure 1A), even though the bacterial cells secreting the inhibitor were genetically identical to the inhibited *Lp*. In both cases, we observed a sharp boundary at the edge of the zone of inhibition. In contrast, the “no zone” *Lp* strain *hisC2*::Tn did not generate a sharp zone of inhibition against neighboring *Lp* (Figure 4A), suggesting that the HGA-melanin pathway was responsible for both *Lm* and *Lp* inhibition. Furthermore, we found that synthetic HGA was able to inhibit *Lp* in liquid cultures (Figure 4B) at the same concentrations that were inhibitory to *Lm* (compare Figure 4B and 3C).

**Figure 4:**
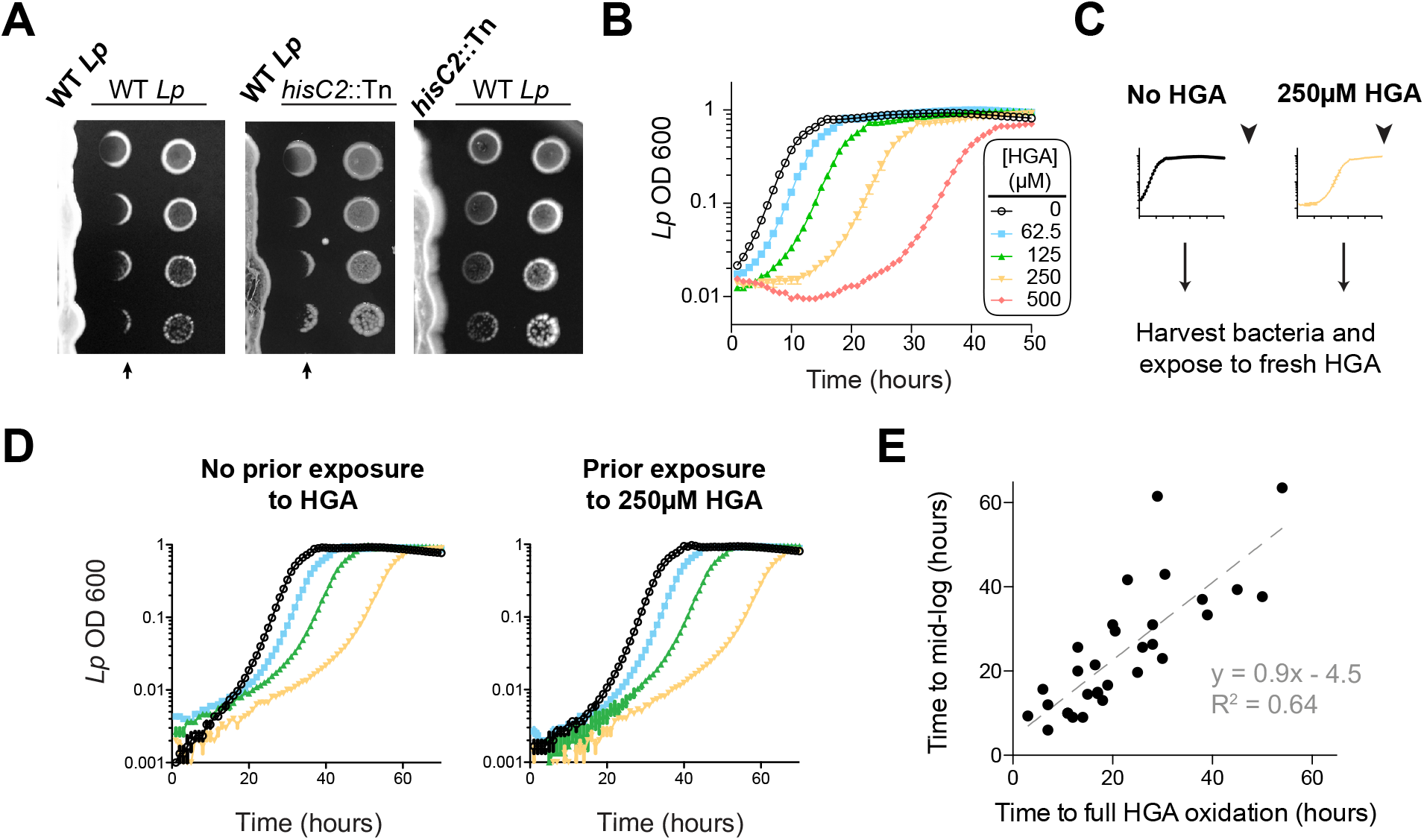
*L. pneumophila* is susceptible inhibition by HGA. **A**) When pre-incubated on agar plates, *Lp* produces a zone of inhibition (arrows), preventing the growth of genetically-identical *Lp* plated 3 days later. The “no zone” mutant *hisC2*::Tn does not produce a sharp front of inhibition, implicating HGA. **B**) Increasing concentrations of synthetic HGA inhibit the growth of *Lp*, causing a growth delay in rich media. Error bars showing standard deviation are small and mostly obscured by the symbols. **C**) To test if Lp population recovery at late time points following HGA exposure is due to the outgrowth of HGA- resistant mutants, we grew *Lp* with or without HGA and sampled bacteria at the end of the experiment (arrowhead) that were unexposed to HGA or were exposed to 250 µM HGA. These were used to inoculate media +/- fresh HGA. **D)** Prior HGA exposure did not lead to subsequent resistance. **E)** The time for *Lp* to grow in the presence of HGA is correlated with the time for synthetic HGA to oxidize at each concentration, suggesting that *Lp* delays growth until HGA has sufficiently oxidized to lose inhibitory activity. Plot shows data combined across 8 experiments. Linear fit curve and equation are shown.

However, our comparisons between *Lm* and *Lp* revealed one important difference in their response to HGA inhibition. Unlike *Lm*, the *Lp* cultures exposed to HGA exhibited a population rebound after a dose-dependent growth delay, measured by both OD600 and plating for viable CFUs (Figure 4B, Supplemental Figure 4C). This rebound response was shared between the KS79 lab strain and the Philadelphia-1 original *Lp* strain (Supplemental Figure 4A). We hypothesized that the *Lp* rebound response following HGA inhibition occurs because of selection and outgrowth of HGA- resistant mutants following exposure. To test this possibility, we collected ‘post-rebound’ stationary phase *Lp* previously exposed to 250µM HGA and compared their subsequent HGA sensitivity to unexposed *Lp* (Figure 4C-D). We found that both cultures showed nearly identical susceptibility to HGA inhibition, suggesting that *Lp* population rebounds are not driven by genetic adaptation.

We therefore considered an alternate possibility that *Lp* populations exposed to HGA remain static until HGA levels fall below inhibitory concentrations, reflecting the auto-oxidation and loss of HGA activity over time. This possibility was supported by CFU measurements, which showed that HGA is bacteriostatic but not bacteriocidal against *Lp* during the growth delay (Supplemental Figure 4C). Furthermore, we observed a strong, linear correlation between the time required to fully oxidize a given concentration of HGA and the length of the growth delay induced by *Lp* (Figure 4E). Based on our multiple observations, we favor the parsimonious conclusions that synthetic HGA causes initial, bacteriostatic inhibition of *Lp* until it has been sufficiently oxidized and thereby inactivated, enabling *Lp* growth. We note that the liquid culture assays (Figure 4B) differ from the co-plating assays, in which we did not observe *Lp* rebound (Figure 4A). In the latter case, we presume that bacteria continually secrete fresh HGA to replace the oxidized HGA over time. Thus, our results confirm a surprising role for HGA in both interspecies and intraspecies *Legionella* inhibition.

### Non-essential gene *lpg1681* sensitizes *L. pneumophila* to HGA

We next investigated the molecular basis of *L. pneumophila* susceptibility and resistance to HGA using bacterial genetics. First, we tested the role of the HmgA-C proteins, which break down intracellular HGA and recycle it back into central metabolism. We hypothesized that HmgA-C proteins might also be able to deactivate extracellular HGA (Rodriguez-Rojas et al. 2009) (Figure 2D). Contrary to this hypothesis, we found that the growth response of the Δ*hmgA* mutant to increasing concentrations of synthetic HGA was nearly identical to that of wild type *Lp* (Supplemental Figure 4B). These results suggest that the intracellular recycling pathway does not play an appreciable role in *Lp* resistance to extracellular HGA.

Having excluded the obvious candidate pathway for *Lp* resistance to HGA, we pursued an unbiased forward genetics approach. Because HGA is strongly inhibitory to low density bacteria, we performed a selection for spontaneous, HGA-resistant mutants of *Lp* and *Lm* using a high HGA concentration that normally prevents almost all growth for both species. To prevent HGA from reacting with media components and becoming inactive (as in Figure 3D, 3F), we mixed the bacteria with 1mM HGA in agar overlays poured onto low-cysteine BCYE plates (Figure 5A). After six days, an average of 53 colonies had grown up on each *Lp* plate under HGA selection, whereas only 3-5 colonies grew on *Lm* plates exposed to with HGA. We also found only 3-5 colonies of either *Lm* or *Lp* exposed to low-cysteine plates without HGA. Based on these results, we focused on HGA-selected mutants of *Lp*. We retested the phenotypes of spontaneous mutants on HGA + low-cysteine plates and recovered 29 *Lp* strains that consistently grew better than wild type *Lp* (Figure 5B). Notably, all recovered mutants had a decrease in HGA sensitivity relative to wild type, but none were completely resistant.

**Figure 5:**
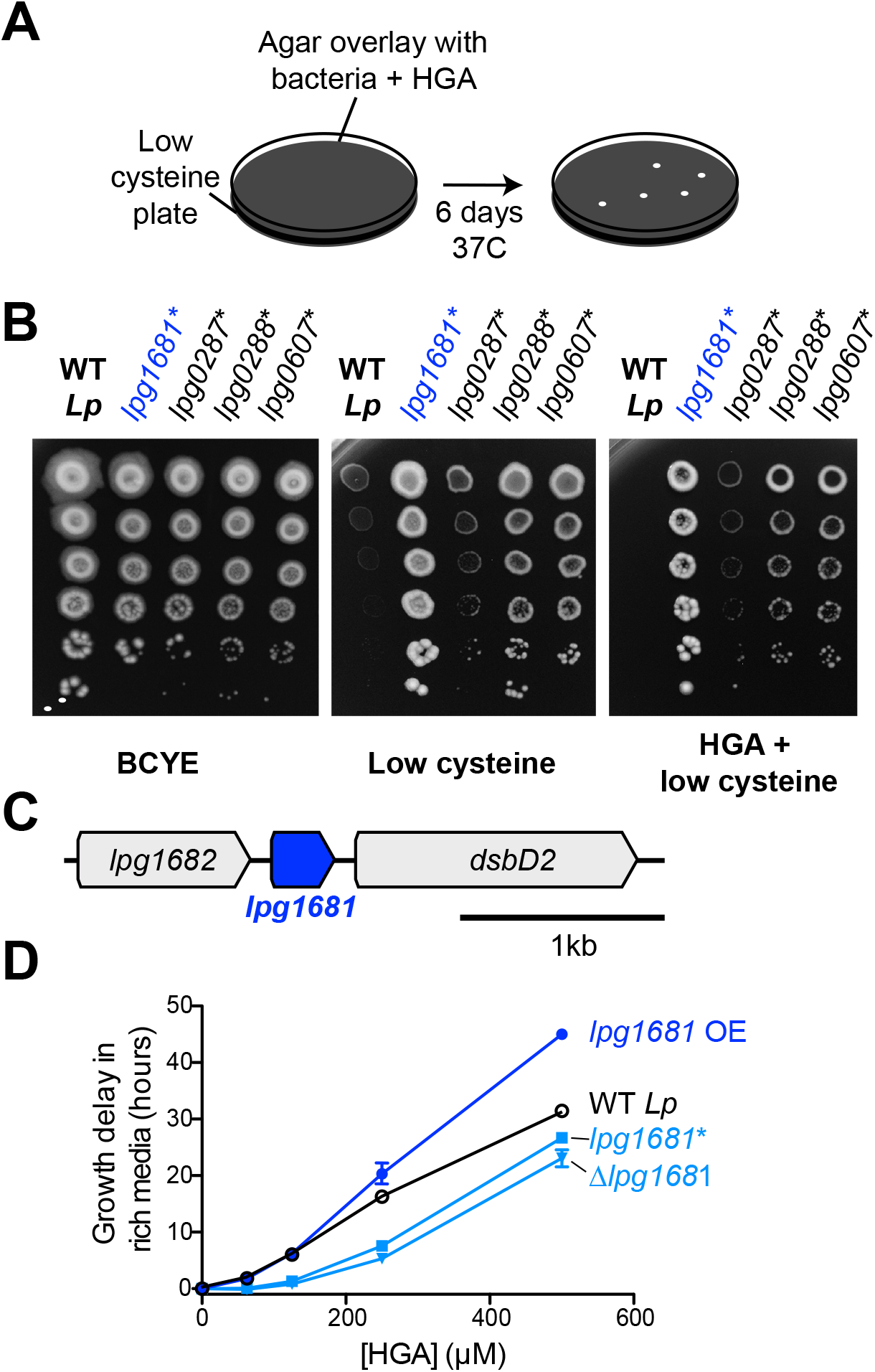
*L. pneumophila* susceptiblity to HGA is modulated by *lpg1681*. **A**) Scheme to select for *Lp* spontaneous HGA-resistant mutants. **B)** Growth of HGA-selected mutants (*) compared to wild type *Lp.* All isolates grew better than wild type in selection conditions (HGA + low cysteine), as well in low cysteine conditions. **C)** Syntenic region of *lpg1681* in *Lp* strains. *lpg1681* is a hypothetical gene that lies downstream of *lpg1682*, a predicted oxidoreductase/dehydrogenase, and upstream of *dsbD2*, a thiol:disulfide interchange protein. **D**) In rich media, a spontaneous *lpg1681* mutant (*lpg1681\**) and the *lpg1681* deletion strain (Δ *lpg1681*) are less sensitive to growth inhibition by HGA than wild type *Lp*, as seen by a shorter growth delay at each concentration of HGA. Overexpression of *lpg1681* (OE) heightens sensitivity to HGA (longer growth delay). Graphs here summarize experiments similar to those in Fig 4B. See Supplemental Figure 8 for full data.

To determine the underlying genetic basis of these phenotypes, we sequenced the genomes of all 29 strains plus the starting, wildtype strain of *Lp* to a median depth of 118x and identified mutations genome-wide. Each mutant strain carried 1 to 3 unique point mutations relative to the starting strain, but most of these mutations were found only in a few shared loci (Table 1). The most abundant category of mutants was genes related to translation. 19 of 29 resistant *Lp* strains carried mutations in translation-related machinery, of which 17 carried mutations in elongation factor P (*lpg0287*) or enzymes responsible for adding post-translational modifications to elongation factor P (*lpg0607* and *lpg0288*). Elongation factor P acts to re-start stalled ribosomes at polyproline tracts, and its post-translational modifications are essential for these functions (Yanagisawa et al. 2010; Navarre et al. 2010; Doerfel et al. 2013; Marman, Mey, and Payne 2014). Based on the frequency of polyprolines in the *Lp* proteome, disruptions to elongation factor P function have the potential to impact the expression of about 33% of *Lp* proteins. The HGA resistance phenotypes we observed in these 17 *Lp* mutants could therefore result from either a large-scale shift in gene expression, or from the altered expression of specific susceptibility genes.

Elongation factor P disruption has also been previously linked to the activation of the stringent response pathway (Nam et al. 2016), which is important for coordinating a variety of stress responses in *Legionella* including oxidative stress (Molofsky and Swanson 2004; Oliva, Sahr, and Buchrieser 2018). We therefore tested if the stringent response pathway is involved in regulating HGA susceptibility or resistance. We assayed HGA susceptibility in mutant *Lp* strains with disruptions to *rpoS*, *letA*, *relA*, and *spoT*, which act early in the stringent response pathway to regulate and/or respond to the levels of the alarmone ppGpp (Supplemental Figure 7C) (Bachman and Swanson 2001; Hammer, Tateda, and Swanson 2002; Dalebroux, Edwards, and Swanson 2009). We found that the HGA susceptibility of all these mutants was similar to that of wild type; furthermore, complementation with these genes did not alter HGA susceptibility (Supplemental Figure 7D). Thus, we find no evidence of a role for the stringent response in HGA susceptibility or resistance. Instead, we propose that disruptions to elongation factor P result in pleiotropic translation defects that together lead to HGA resistance via still-unknown mechanisms. In addition to the translation-related mutants, we found 5 missense mutations in *secY* or *secD* (*lpg0349* and *lpg2001*), members of the Sec secretion apparatus that moves polypeptides across the cytosolic membrane; 1 mutation in *aceE* pyruvate dehydrogenase (*lpg1504*); and 4 mutations in a hypothetical gene, *lpg1681* (Table 1, Figure 5C). The Sec apparatus is involved in the secretion of many substrates and mutations to this machinery could also lead to extensive pleiotropic defects.

Instead, we focused on the relatively uncharacterized hypothetical gene *lpg1681*, which encodes a small, 105 amino acid protein with no predicted domains apart from two transmembrane helices. This gene is adjacent in the genome to *lpg1682*, which encodes for a predicted oxidoreductase/dehydrogenase, and *lpg1680*, which encodes for the thiol:disulfide exchange protein DsbD2 (Figure 5C, Supplemental Figure 6). Functional studies of DsbD2 (aka DiSulfide Bond reductase D2) have demonstrated that it interacts with thioredoxin to regulate disulfide bond remodeling in the periplasm (Inaba 2009; Kpadeh et al. 2015). If *lpg1681* has a redox function related to its neighboring genes, we expected its syntenic locus to be conserved across bacterial strains and species. Consistent with this prediction, we found that the *lpg1680*-*1682* locus is present and conserved among over 500 sequenced *Lp* strains currently in NCBI databases. Outside *L. pneumophila*, *lpg1681* is mostly restricted to the *Legionella* genus, present in about half of the currently sequenced species (Burstein et al. 2016) (Supplemental Figure 6). A homolog of *lpg1681* is also found in the draft genome of *Piscirickettsia litoralis*, a gamma proteobacterium and fish pathogen (Wan et al. 2016). In all cases, *lpg1681* resides upstream of *dsbD2*, suggesting a functional link between these proteins (Supplemental Figure 6) and implicating *lpg1681* in a role in redox homeostasis. We, therefore, viewed *lpg1681* as a promising candidate for a gene involved in HGA susceptibility.

We constructed *lpg1681* overexpression and deletion strains in *Lp* and tested the susceptibility of these strains to HGA. Similar to the spontaneous mutants we recovered, we found that the Δ*lpg1681* strain was more resistant to synthetic HGA in rich media (Figure 5D, Supplemental Figure 5). Conversely, overexpression of *lpg1681* increased *Lp* sensitivity, resulting in longer growth delays than wild type at high concentrations of HGA. We therefore conclude that wild type *lpg1681* expression sensitizes *Lp* to inhibition by extracellular HGA. Given its genetic linkage with *DsbD*, our findings further suggest that alteration of disulfide bond regulation in the periplasm might constitute one means to mitigate HGA susceptibility.

### *L. pneumophila* susceptibility to HGA is density-dependent

Our unbiased genetic screen for *Lp* resistance to HGA only revealed mutants that were partially resistant to HGA. These mutants had a smaller growth delay than wild type at a given HGA concentration, but all remained qualitatively susceptible to inhibition. These results suggest that the genetic routes for *Lp* to completely escape from HGA-mediated inhibition are limited. Yet, *Lp* secretes abundant HGA into its local environment, despite the fact that HGA secretion is not required for *Lp* growth or metabolism (as seen by the robust growth of the *hisC2::Tn* mutants, Figures 2 and 4). Thus, our findings do not provide an adequate explanation for the paradox of how *Lp* cells can secrete a toxic compound to which they apparently carry no heritable resistance.

We therefore considered a distinct mechanism by which *Lp* might avoid self-inhibition: *Lp* might produce and secrete HGA only during conditions to when it is not susceptible to HGA. To address this possibility, we investigated when and where *Lp* secretes HGA. We tracked HGA secretion across a growth curve of *Lp* in rich media, using our previously described assay of using a standard curve of synthetic HGA to estimate HGA levels. It has long been known that *Lp* produces abundant HGA-melanin pigment in stationary phase, when the bacteria are undergoing very slow or no growth (Pine et al. 1979; Berg et al. 1985; Wiater, Sadosky, and Shuman 1994). By comparing to a synthetic HGA standard curve (Supplemental Figure 4F), we estimate that *Lp* produces a burst of HGA in stationary phase, secreting over 250µM within 5 hours (Figure 6A). HGA secretion then continues after the population has ceased growing. These quantities of HGA are more than enough to be inhibitory to *Lp* growth (Figure 4B).

**Figure 6:**
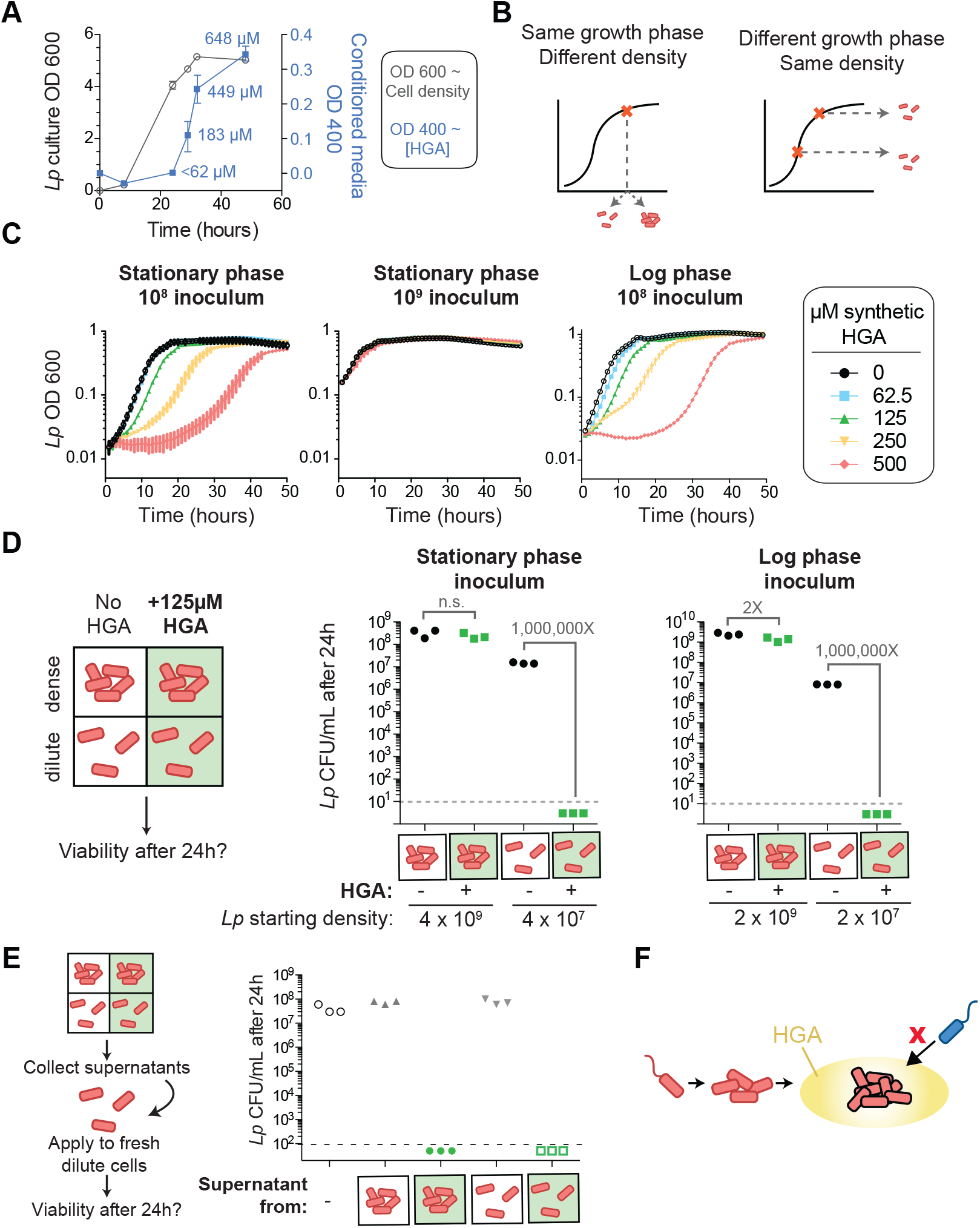
HGA susceptibility in *Lp* is linked to cell density, independent of inoculum growth phase**. A)** Timing of HGA secretion by *Lp* in rich media, measured by OD400 of conditioned media (CM) after allowing for full oxidation (blue boxes, right y-axis). A matched growth curve of *Lp* is presented for comparison (gray circles, left y-axis). Abundant HGA is secreted in during stationary phase. Estimates of secreted HGA concentration (blue) are based on a standard curve of synthetic HGA. Error bars show standard deviations. **B)** Schematic showing how experiments controlled for inoculum density vs. growth phase. To test the impact of cell density, a single culture was diulted to multiple densities at the start of the experiment. To test the impact of growth phase, *Lp* was sampled at multiple stages of growth and diluted to the same CFU/mL at the start of the experiment. **C**) *Lp* growth is inhibited by HGA when cells are inoculated at relatively low density (10^8 CFU/mL), but HGA is ineffective in inhibiting growth when cells are inoculated at high density (10^9 CFU/mL). Growth phase of *Lp* inoculum has little impact on HGA susceptibility. **D**) Viable CFUs following 24h incubation of *Lp* with or without 125µM HGA in nutrient-free PBS. When incubated at high density, bacteria are mostly resistant to HGA, while they are higly sensitive at lower density. Dotted line shows limit of detection. Brackets indicate the fold change in viable CFUs due to HGA exposure **E**) High-density cells do not inactivate extracellular HGA. Supernatants from *Lp* +/- HGA at two densities were collected and applied to dilute cells for 24h. Viable CFUs were counted following supernatant exposure. HGA in all supernatants remained active against low density *Lp*. **F)** Model for HGA activity. *Lp* (orange) colonizes a surface and grows to form a microcolony. Once cells are at high density, they secrete abundant HGA (yellow). Through unknown mechanisms, high density *Lp* are resistant to HGA’s effects (black outline), while low density *Lp* or other *Legionella* species (blue) are inhibited by HGA and cannot invade the microcolony’s niche.

For *Lp* to avoid self-inhibition from HGA, we hypothesized that *Lp* might be resistant to this inhibitor at high density and/or when the cells are in a stationary phase of growth. We therefore investigated if cell density or growth phase impacted HGA susceptibility (Figure 6B). For *Lp* exposed to HGA in rich media, we found that the growth phase of bacteria used to inoculate the experiment had little impact on HGA susceptibility (Figure 6C). In contrast, when cells were inoculated at high density (10^9/mL instead of 10^8/mL), they were resistant even to high concentrations of HGA (Figure 6C), suggesting that cell density might be linked to HGA resistance. However, because both cell density and growth phase are changing over time during our assays in rich media, we cannot fully separate their contributions to HGA-resistance in *Lp*. We therefore created a new assay to assess HGA susceptibility. We exposed *Lp* bacteria to HGA at different dilutions in nutrient-free PBS, which ensured that the bacteria did not grow or change cell density during the course of the experiment. After 24 hours exposure to 125 µM HGA, we assessed bacterial viability by plating for viable CFUs (Figure 6D). No measurable darkening of the HGA was detected in this assay, suggesting that the oxidation and de-activation of HGA was considerably slowed in low-nutrient conditions. We found that *Lp* bacteria incubated at high density with HGA (above 10^8 CFU/mL) were largely protected from inhibition (Figure 6D). However, at lower density (10^7 CFU/mL), *Lp* bacteria were extremely sensitive to HGA, with at least a 10^6-fold reduction in CFUs, to below our limit of detection. As in the rich media assay, this density-dependent resistance to HGA was not altered by inoculum growth phase (Figure 6D).

We reasoned that the quorum sensing pathway was most likely to control the substantial, density-dependent difference in HGA susceptibility. We therefore investigated if HGA resistance depended on the previously described *Lp* quorum sensing response regulator, *lqsR* (Tiaden et al. 2007). We found that deleting *lqsR* had no detectable impact on HGA susceptibility or resistance (Supplemental figure 7B). Therefore, the density-dependent susceptibility of *Lp* to HGA must be independent of the *lqsR* pathway.

We next investigated the basis of high-density resistance to HGA by *Lp* bacteria. We considered the hypotheses that high-density cells could alter the activity of extracellular HGA, either through the secretion of inactivating compounds or through bulk, non-specific binding of HGA to bacterial biomass, leading to a reduction in its effective concentration. To test both hypotheses, we recovered the supernatants from high-density and low-density bacteria that had been incubated with or without HGA, and applied these supernatants to fresh, low-density *Lp* to test their HGA activity (Figure 6E). We found that the supernatants from HGA-exposed *Lp* remained fully inhibitory, even after 24h incubation with high-density bacteria. Furthermore, we found that adding heat-killed *Lp* bacterial cells to low-density viable *Lp* bacteria did not enhance the latter’s resistance to HGA inhibition (Supplemental Figure 7A). Therefore, we conclude that HGA susceptibility appears to be density-dependent and yet cell-intrinsic. Because *Lp* bacteria at high density both secrete and are protected from HGA, this strategy of secreting HGA only when *Lp* cells are conditionally HGA-resistant may allow *Lp* to produce a broadly active inhibitor while restricting the potential for self-harm.

## Discussion

The HGA-melanin pathway is well-studied and widespread among bacteria and eukaryotes. In this study, we identify HGA as a mediator of inter-*Legionella* inhibition, both between *Legionella* species and even between genetically identical populations of *L. pneumophila*. To our knowledge, this is the first time that HGA has been described to have antimicrobial activity. One reason HGA-mediated inhibition may not have been previously documented is that HGA is a redox-active, unstable molecule with transient activity. Our study finds that synthetic HGA can auto-oxidize over the course of an experiment to form inactive HGA-melanin (Figure 3, Supplemental Figure 4D), allowing exposed *Lp* populations to rebound following initial inhibition (Figure 4B). Intriguingly, although *Lp* populations recover upon HGA oxidation, *Lm* populations do not, suggesting that HGA may cause more harm to *Lm* cells. We find that the quenching of HGA’s inhibitory activity occurs even more rapidly in the cysteine-rich microbial media typically used to grow *Legionella* in the lab (Supplemental Figure 4D). Conversely, we find that HGA becomes more potent in low-cysteine media or in PBS (Figures 3F, 6D), where oxidation of HGA into HGA-melanin occurs more slowly; such nutrient-poor conditions may better replicate the nutrient-poor (oligotrophic) aquatic environments of *Legionella’s* natural habitat (Atlas 1999; Boamah et al. 2017).

Although dense, stationary-phase cultures of *L. pneumophila* secrete abundant HGA (Figure 6A, Supplemental Figure 2D), we also find that these bacteria do not possess heritable resistance to HGA (Figure 4B-D) and are highly susceptible to HGA inhibition at low cell density. However, they exhibit conditional, density-dependent, cell intrinsic resistance to HGA at high cell density (Figure 6). This lack of heritable resistance makes HGA-mediated inhibition different from classical antibiotics or toxins, which typically are produced by bacteria that also express resistance genes or antitoxins. HGA inhibition is also distinct from that caused by toxic metabolic by-products in two important ways. First, HGA production is not required for efficient growth or metabolism in *Legionella* species (Supplemental Figure 2D, and see growth of *hisC2*::Tn mutant in Figure 4A). Second, high-density populations of *L. pneumophila* that produce HGA are themselves protected from by HGA inhibition (Figure 6). Only neighboring, low-density *Legionella* are strongly inhibited (Figure 4A, 6D). The strong density-dependence of *Legionella*’s susceptibility to HGA may be another reason that it has been previously undiscovered despite intense study of these bacteria.

HGA-mediated inhibition of *Legionella* is reminiscent of the antimicrobial activities of phenazines, another class of small aromatic molecules such as pyocyanin from *Pseudomonas aeruginosa*. Both types of molecules are redox-active (Hassan and Fridovich 1980), are produced at high cell density (Hassan and Fridovich 1980; Baron, Terranova, and Rowe 1989), are able to chemically react with thiol groups (Cheluvappa et al. 2008; Heine et al. 2016), and result in the production of a colored pigment (Price-Whelan, Dietrich, and Newman 2006). Phenazine inhibitory activity typically comes from redox cycling and the production of reactive oxygen species, including H_2_O_2_ (Hassan and Fridovich 1980; Cheluvappa et al. 2008). Oddly, HGA-melanin production has previously been implicated both in the production of (Noorian et al. 2017) and protection from (Keith et al. 2007; Orlandi et al. 2015) reactive oxygen species. The catalase experiments presented here have ruled out the production of extracellular H_2_O_2_ as a possible mechanism explaining HGA inhibition (Supplemental Figure 3B-C). Instead, based on association of *lpg1681* and *DsbD2* with reduced HGA sensitivity (Figure 5), the ability of diverse thiols to quench HGA’s activity (Supplemental Figure 3E), and precedents from phenazines (Heine et al. 2016), we propose that HGA (and/or its transient, oxidative intermediates) may be toxic by forming adducts on cysteine residues or otherwise disrupting disulfide bonding. Alternatively, HGA inhibition could occur via the production of other reactive oxygen species, including potentially the generation of intracellular superoxide and/or H_2_O_2_ (Hassan and Fridovich 1979), which would not be affected by catalase treatment. As both of these mechanisms should be inhibitory to a number of bacterial taxa, it will be interesting to broadly survey HGA susceptibility outside of *Legionella*.

The density-dependence of *Lp*’s resistance to HGA is unusual and worthy of future study. Because high-density cells do not inactivate or bind up extracellular HGA (Figure 6E) and because heat-killed cells cannot protect live, low-density *Lp* from inhibition (Supplemental Figure 7A), we infer that resistance is cell-intrinsic, resulting from differing physiology and/or gene expression between high-and low-density cells. Two pathways that commonly regulate such defenses include the stringent response pathway, which becomes active under nutrient limitation and stress, and quorum sensing pathways, which become active at high cell density (Bachman and Swanson 2001; Hammer, Tateda, and Swanson 2002; Dalebroux, Edwards, and Swanson 2009; Hochstrasser and Hilbi 2017). Although our experiments disrupting these pathways suggest that neither pathway contributes to density-dependent susceptibility or resistance (Supplemental Figure 7B-D), we note that quorum sensing in *Legionella* remains understudied, and such pathways vary considerably across bacterial taxa (Hochstrasser and Hilbi 2017; Miller and Bassler 2001). Future work using unbiased approaches to investigate the regulation of HGA susceptibility may be able to uncover additional density-sensing pathways, possibly including an undescribed mode of quorum sensing in *Legionella*.

*L. pneumophila* is often co-isolated with other *Legionella* species, which likely compete for similar ecological niches (Wery et al. 2008; Pereira et al. 2017). HGA- mediated inter-*Legionella* inhibition therefore has a strong potential to impact the success of *Lp* in both natural and manmade environments. Because high-density, established *Lp* bacterial communities are largely resistant to HGA inhibition, these communities might use HGA to protect against low-density, invading *Legionella* competitors with little harm to themselves (Figure 6F). In this model, motile *Lp* can disperse, colonize a new surface, and grow into a microcolony using the locally available nutrients. In these early stages, no HGA is produced. After the *Lp* population grows up and crosses a certain cell density threshold, the cells become HGA-resistant through a cell-intrinsic mechanism. When this dense population enters stationary phase, it also begins to secrete abundant HGA into the local environment. This secreted HGA has minimal impact on the resistant, producer cells. However, it can inhibit the growth of nearby, low-density *Legionella*, whether the neighboring cells are other *Legionella* species or even genetically-identical *Lp*. Given these dynamics observed in the lab, we speculate that HGA and other such inhibitors may be deployed as a bacterial niche-protective strategy.

## Materials and Methods

### Bacterial strains and growth conditions

The bacterial strains and plasmids used for this study are listed in Supplemental Table 2. As our wild type *Legionella pneumophila (Lp)* strain, we used KS79, which was derived from JR32 and ultimately from isolate Philadelphia-1 (de Felipe et al. 2008; Sadosky, Wiater, and Shuman 1993; Rao, Benhabib, and Ensminger 2013). Compared to JR32, the KS79 strain has a *comR* deletion to enable genetic manipulation (de Felipe et al. 2008). We used *Legionella micdadei (Lm)* tatlock as our susceptible strain (Garrity, Brown, and Vickers 1980; Hébert, Steigerwalt, and Brenner 1980). Liquid cultures of *Legionella* were grown shaking in AYE rich liquid media at 37°C (De Jesús, O’Connor, and Isberg 2013). Unless otherwise indicated, experiments were inoculated with stationary phase *Legionella*, grown from a single colony in AYE for 16-18 hours as described, to a density of 3 − 4 × 10^9 CFU/mL. For experiments with log phase *Lp*, an overnight culture was diluted into fresh AYE at 1:8 ratio (to a density of 4 − 5 × 10^8 CFU/mL) and allowed to grow to a density of 10^9 CFU/mL before setting up the experiment.

To manipulate the redox state of AYE, we altered the amount of cysteine added to the media from 0.4 g/L in standard AYE to 0.1, 0.2, and 0.8 g/L. On solid media, *Legionella* were grown either on BCYE agar plates either containing the standard cysteine concentration (0.4g/L) or in “low cysteine” conditions (0.05g/L) (Feeley et al. 1979; Solomon and Isberg 2000). *E. coli* strains used for cloning were grown in LB media. Where indicated, antibiotics were used at the following concentrations in solid and liquid media, respectively; chloramphenicol (5 µg/mL and 2.5 µg/mL), kanamycin (40 µg/mL) and ampicillin (50 µg/mL and 25 µg/mL). For counter-selection steps while generating deletion strains, 5% sucrose was added to BCYE plates. For agar overlay experiments, we used 0.7% agar dissolved in water, which was kept liquid at 50°C before pouring over low cysteine BCYE plates.

### Gene deletions and complementations

Genomic knockouts in *L. pneumophila* were generated as previously described (Wiater, Sadosky, and Shuman 1994). Briefly, we used an allelic exchange plasmid (pLAW344) harboring chloramphenicol and ampicillin selection cassettes and the counter-selection marker SacB, which confers sensitivity to sucrose. Into this plasmid, we cloned ∼1 kb regions upstream and downstream of the gene of interest to enable homologous recombination. Following electroporation and selection on chloramphenicol, we used colony PCR to verify insertion of the plasmid into the chromosome, before counter-selection on sucrose media. From the resulting colonies, we performed PCR and Sanger sequencing to verify clean gene deletion. For complementation, the coding region of a candidate gene was cloned into a plasmid (pMMB207c) following a ptac promoter (J. Chen et al. 2004). To induce gene expression, strains carrying pMMB207c-derived plasmids were exposed to 1mM IPTG. All constructs were assembled using Gibson cloning (NEB Catalog #E2621).

### Inhibition assays on agar plates

To visualize inhibition between neighboring *Legionella* on solid media, a streak of approximately 5 × 10^6 CFU of the inhibitory strain of *Lp* was plated across the center of a low cysteine BCYE plate. After 3 days growth at 37°C, dilutions of susceptible *Lp* or *Lm* were plated as 10µLspots approximately 1 cm and >2 cm from the central line. Once spots were dry, plates were then incubated for an additional 3 days before scoring for inhibition. This assay was also used to quantify the bactericidal inhibition of *Lm*, with slight modifications. Here, all *Lm* was plated in 20µL spots at 10^6 CFU/mL. The time of plating susceptible *Lm* was treated as t=0. Once spots were dry, plugs were extracted from within the *Lm* spots using the narrow end of a Pasteur pipette. These plugs were transferred into media, vortexed, and plated to quantify CFU. This procedure was repeated after 48h at 37°C to compare *Lm* viability and growth within (“near”, Figure 1B-C) or outside (“far”) of the zone of inhibition.

For inhibition assays on bacterial lawns, we plated 10µL drops of either live *Lp* or chemical compounds on top of a lawn of 5 × 10^7 CFU *Lm* on low cysteine BCYE, and assessed growth of the lawn after 3 days at 37°C. Synthetic HGA (Sigma: #H0751) was dissolved in water at a concentration of 100mM and filter sterilized before use. To limit the potential for HGA oxidation prior to use, 100mM aliquots prepared in water were stored frozen at −20C and discarded after 1-2 weeks. HGA-related compounds, 2- hydroxyphenylacetic acid (Sigma: #H49804) and 3-hydroxyphenylacetic acid (Sigma: #H49901), were prepared in the same way. To test the impact of DTT (Sigma: #43819) and glutathione (oxidized: Sigma #G4376, reduced: Sigma #G6529) on HGA-mediated inhibition, filter-sterilized solutions dissolved in water were mixed in equimolar ratios with HGA, and incubated shaking at room temperature for 1 hour before spotting onto bacterial lawns. HGA-melanin pigment was prepared from *Lp* conditioned media as previously described (Zheng et al. 2013) from KS79, the unpigmented *hisC2*::Tn mutant, and the hyperpigmented Δ*hmgA* mutant. Briefly, conditioned media was collected and sterile filtered from 100 mL cultures of *Lp* in AYE media grown shaking at 37°C for 3 days. The conditioned media was acidified to a pH of 1.5 and transferred to 4°C for 2 hours to precipitate. Precipitated pigment was collected by centrifugation at 4000 × g for 15 minutes and then washed with 10mM HCl. Pelleted pigment was then returned to neutral pH and resuspended in PBS at 10X before testing.

### Transposon mutagenesis screen

For random transposon insertion mutagenesis, we used a Mariner transposon from the pTO100 plasmid (O’Connor et al. 2011). We electroporated this plasmid into the KS79 strain and allowed cells to recover at 37°C for 5 hours. To select for cells with integrated transposons, cultures were plated on BCYE/Kan/sucrose plates and incubated at 37°C for 3 days before screening individual mutant colonies.

To identify transposon mutants with defects in *Lm* inhibition, we transferred each *Lp* mutant onto a low cysteine plate with a lawn of 5 × 10^7 CFU *Lm* and visually screened for those with either small zones of inhibition or no zone of inhibition. This transfer of *Lp* mutants was achieved either by replica plating using sterile Whatman paper (Whatman: #1001150) or by manual transfer with a sterile toothpick. Plates were then incubated at 37°C for 3 days and scored. All putative mutants underwent clonal re-isolation, were diluted to OD 600 of 0.1, and spotted on fresh *Lm* lawns to retest their phenotypes. To map the sites of transposon integration, we used arbitrary PCR as described in (T. Chen et al. 1999), with primers redesigned to work with the pTO100 transposon (Table 3). Briefly, this protocol involved two PCR steps to amplify the DNA flanking the transposon. The first step used low annealing temperatures to allow the arb1 randomized primer to bind many sites in the flanking DNA while the pTO100_F or pTO100_R primer annealed within the transposon, generating multiple products that overlapped the flanking DNA. These products were amplified in the second step PCR using the arb2 and pTO100_Rd2 primers, and we used the pTO100_Rd2 primer for Sanger sequencing. PCR programs and conditions were as in (T. Chen et al. 1999).

### HGA inhibition assays in AYE rich media

For rich media assays (e.g. Figures 3C-F, 4B, 4D), overnight cultures of *Legionella* were diluted to 10^8 CFU/mL in AYE, mixed with synthetic HGA (at 0, 62.5, 125, 250, or 500 µM final) or with isolated pigment in 96 well plates, and grown shaking at 425 cpm at 37°C. The cytation 3 imaging reader (BioTek™ CYT3MV) was used to monitor growth by OD 600 measurements. Because oxidized pigment from synthetic HGA is detected at OD 600 as well, each experiment included bacteria-free control wells containing media and each concentration of HGA. To correct OD 600 readings for pigment development, at each time point we subtracted the control well reading from bacterial wells that received the same concentration of synthetic HGA.

For experiments with HGA “pre-oxidation” (Figure 3D), we diluted HGA in AYE media and incubated this solution shaking at 37°C for 24 hours in the plate reader before adding *Lm* bacteria. Complete oxidation of HGA during the 24 hours was monitored using OD400 to track the accumulation of HGA-melanin pigment (Supplemental figure 4). To test if extracellular catalase could protect from HGA inhibition (Supplemental figure 3B), we incubated *Lm* at 10^7 CFU/mL with or without 125 µM HGA and either 0, 1, 10, or 100 U/mL of bovine catalase (Sigma #C30). As a control to ensure that the catalase was active, we incubated the bacteria as above with catalase and 2mM H_2_O_2_ (Sigma #88597).

In *Lp*, HGA inhibition in AYE rich media resulted in a growth delay, similar to an extended lag phase (Figure 4B). To determine if this delay was due to genetic adaptation, we sampled *Lp* after 70 hours growth with 250 µM HGA or without HGA (Figure 4C-D). These bacteria were washed once and resuspended in fresh AYE, before being diluted back to 10^8 CFU/mL and then exposed to fresh, synthetic HGA as above. To assess the correlation between HGA oxidation and the length of the *Lp* growth delay (Figure 4E), we pooled data from 8 experiments on different days that measured wild type *Lp* (KS79) exposed to a range of HGA concentrations in AYE. We considered the “Time to HGA saturation” as the length of time required for a given concentration of HGA to stop forming additional HGA-melanin, measured as the time until OD400 readings increased by less than or equal to 0.001 units per hour. The “Time to mid-log” was measured as the time when *Lp* exposed to that concentration of HGA had grown to an OD600 of 0.1. To compare sensitivity to HGA among *Lp* strains, we calculated the lag phase from the growth curve of each well using the GrowthRates program (Hall et al. 2014). We excluded a small number of samples where the growth curve was not well fit (R<0.99), and then for each strain used the difference in lag time between the samples with and without HGA to calculate the growth delay due to HGA (Figures 5C).

### HGA inhibition assays in PBS

While we were able to manipulate inoculum growth phase and cell density in the AYE assays, during these experiments the bacteria altered their density and growth phase as they grew in rich media. To separate the impacts of cell density and growth phase on HGA susceptibility, we used a complementary assay in which we evaluated *Legionella* viability when exposed to HGA in nutrient-free PBS at different cell densities. This design ensured that the bacteria maintained a constant cell density throughout the course of the experiment. Stationary phase cultures were washed once and re-suspended in PBS. We diluted these bacteria to estimated starting concentrations of 10^9, 10^8, and 10^7 CFU/mL and plated for CFU at t=0. We distributed the remaining bacteria into 96 well plates with or without 125 µM HGA. Plates were incubated shaking in a plate reader at 425 cpm at 37°C for 24 hours before plating to quantify CFU on BCYE plates. CFU were counted after 3-4 days growth at 37°C.

To determine if high density bacteria were protected via mass action effects that diluted out the amount of HGA per cell through binding of bulk material, we asked if the addition of dense, heat-killed bacteria could protect low-density *Lp* from HGA (Supplemental figure 7A). To prepare high-density heat-killed bacteria, an overnight culture of *Lp* was washed once in PBS, resuspended to 2 × 10^9 CFU/mL, and incubated at 100-110 °C for 60 minutes. After heating, this sample was diluted 1:2 and mixed with 10^7 CFU/mL live *Lp* in PBS +/- 125µM HGA to assess protection. As a control, 10^9 and 10^7 CFU/mL live *Lp* with or without HGA were tested simultaneously. Cells were incubated and plated as above to assess viability. To determine if high density bacteria were protected via HGA degradation, we tested if the supernatants from HGA-exposed, high-density bacteria retained the potency to inhibit low-density bacteria (Figure 6E). To generate supernatants, we set up 2mL samples containing 10^8 or 10^7 CFU/mL of *Lp* in PBS +/- 125µM HGA and incubated them shaking at 37°C for 20 hours. After plating aliquots for viable CFU, we pelleted the remaining bacteria and sterile filtered 1mL of each supernatant through a 0.2µm filter. Each supernatant was tested in triplicate, incubated with fresh *Lp* at 10^7 CFU/mL in a 96 well plate as above for 24 hours before plating for CFU. As a control, 10^7 CFU/mL live *Lp* were incubated in PBS alone.

### Quantification of HGA’s ability to react with cysteine

HGA is known to be a redox-active molecule, with a redox potential of +0.636 V (Eslami, Namazian, and Zare 2013). As this measurement can be altered by pH and temperature, we assessed the ability for HGA to oxidize cysteine in our experimental conditions using Ellman’s reagent (also known as 5,5’-dithiobis-(2-nitrobenzoic acid), DTNB, Invitrogen #D8451). Ellman’s reagent reacts in the presence of reduced thiol groups on L-cysteine to form a yellow color, which can be read as 412nm absorbance. When thiol groups are oxidized, the Ellman’s reagent is colorless. We used the ability for HGA to decrease the amount of reduced cysteine as a proxy for its oxidizing ability. Stock solutions of both 100mM HGA and 1.5mM L-cysteine (Sigma #C6852) were prepared fresh in PBS at the start of the experiment. Different concentrations of HGA (from 125µM to 8mM) were incubated in triplicate, shaking at 25°C with 1.5mM cysteine in PBS for 16 hours. These conditions were compared to a standard curve of cysteine from 0-2mM, which were incubated in parallel with the experimental samples to account for cysteine oxidation over time. To quantify the remaining free thiol groups, 180uL of 0.08 mg/mL Ellman’s reagent was mixed with 17.65uL of each experimental or standard sample in a 96 well plate, incubated for 3 hours at 25°C, and read for 412 nm absorbance. The “decrease in reduced cysteine” was calculated as the difference between the initial and final measured cysteine concentrations, based on the standard curve conversion.

### Estimation of amount of HGA secreted by *Lp*

HGA-melanin is a black-brown pigment that is easily detected at OD 400. We took advantage of this coloration to estimate the amount of HGA that had been secreted by *Lp* by comparing the color of conditioned media to a standard curve of oxidized synthetic HGA. To isolate conditioned media from pigment mutant strains, cultures of KS79, Δ*hmgA*, and HisC::Tn were inoculated with fresh colonies from a BCYE plate into 5 mL AYE and were grown shaking at 37°C for 48 hours. We then collected conditioned media by pelleting the bacteria and passing the supernatant through a 0.2µm filter. To harvest conditioned media for a time course, cultures of *Lp* were inoculated into 5 ml AYE and grown shaking at 37°C. After 15, 20, 24, 39, 44, and 48 hours, we measured the OD 600 of the culture and collected conditioned media. To create a standard curve, we diluted synthetic HGA into AYE at the following concentrations: 62.5 µM, 125 µM, 250 µM, 500 µM, and 1mM. The conditioned media and standard curve samples were incubated in a 96 well plate in a plate reader shaking at 37°C for 24 hours to allow the HGA to oxidize. We used OD 400 data from the 24 hour time point to generate a standard curve for each HGA concentration and calculated a line of best fit using linear regression. This equation was used to estimate the amount of secreted HGA that corresponded to the OD 400 of each conditioned media sample. (Supplemental Figure 4).

### HGA-resistant mutants

Because the inhibitory activity of HGA is quenched through interactions with cysteine in rich media (e.g. Figure 3F), it was not possible to select for HGA-resistant mutants by mixing HGA into BCYE agar. Instead, to reduce the potential for HGA to react with media components while allowing sufficient access to nutrients for mutant cells to grow, we selected for HGA-resistant mutants by mixing 4 × 10^7 CFU *Legionella* with 1mM HGA in 4mL of 0.7% molten agar and pouring this solution as an overlay on a low cysteine BCYE plate. Plates were incubated at 37°C for 6 days, before candidate resistant colonies were picked and clonally isolated. The HGA resistance and growth of each isolate was re-tested on overlays with or without 1mM HGA on both regular and low cysteine BCYE.

Twenty-nine isolates were more resistant to HGA than wild type *Lp* upon retesting. We sequenced and analyzed genomic DNA from these isolates and a matched wild type strain as follows. DNA was prepared from each strain using a Purelink genomic DNA mini kit (Invitrogen, #K1820). DNA concentrations were quantified using Qubit and normalized to 0.5 ng/uL. Barcoded libraries were prepared using tagmentation according to Baym et al. 2015 (Baym et al. 2015), analyzed with QuantIT DNA quantification, pooled, and sequenced with 50 bp paired-end reads on an Illumina HiSeq 2500 machine. Reads were trimmed for quality and to remove Nextera indices with Trimmomatic (Bolger, Lohse, and Usadel 2014) and mapped to the Philadelphia-1 genome (Chien et al. 2004) using Bowtie2 with default parameters (Langmead et al. 2009). Coverage plots were generated for each strain using bamcoverage (Ramírez et al. 2016) and manually examined for evidence of large genomic deletions and amplifications. None were observed, apart from a prophage that was present in the reference genome but missing from all sequenced strains, including our wild type KS79 strain. Variants were detected for each mutant using Naive Variant Caller (Blankenberg D, et al. In preparation). Those variants that were detected in mutant strains but not the wild type strain were considered as putative causative mutations. For each of these mutations, we inspected the mapped reads and excluded faulty variant calls that either were adjacent to coverage gaps or that did not appear to be fixed in the clonal mutant and/or wild type sequences, likely due to errors in read mapping. After this manual filtering, 1-3 well-supported mutations remained for each mutant genome. Nine of the mutants were isolated on a different day from the other mutants: in addition to various unshared mutations, these nine strains each carried exactly the same missense mutation in *rplX*, which we disregarded as a background mutation that likely arose before selection. Following this exclusion, each mutant carried only a single well-supported mutation in a coding region. Most often this coding mutation was the only mutation we detected, although one mutant carried two additional intergenic point mutations. The coding mutations were point mutations or small deletions that resulted in non-synonymous changes, frame shifts, or gene extensions. Across different mutants, the mutations we uncovered were repeatedly found in the same, few loci (Table 1).

### Evolution of *lpg1681*

The genes in the HGA-melanin synthesis pathway are highly conserved in diverse bacteria and across the *Legionella* genus, with all genes present in all 41 currently sequenced *Legionella spp.* genomes (Burstein et al. 2016). In contrast, we were able to identify *lpg1681* in only 30 *Legionella spp.* genomes, as well as a single draft genome outside the *Legionella* genus– *Piscirickettsia litoralis*, an intracellular fish pathogen (Wan et al. 2016). Across the *lpg1681*-containing genomes, there is evidence for extensive recombination of the flanking loci, yet *lpg1681* is always found upstream of *dsbD2*. We identified most of these homologs of *lpg1681* using a jackhmmr search (Finn et al. 2015), followed by cross-referencing the homologs with the *Legionella* orthology groups defined by Burstein et al., 2016. From this starting set, additional *lpg1681* orthologs were identified in unannotated, intergenic regions by searching for >200bp open reading frames upstream of *dsbD* orthologs, and confirming the homology of these regions using MAFFT alignments (Katoh et al. 2002). Through this method, we located *lpg1681* in all currently sequenced *Legionella* genomes that contain an annotated *dsbD2* gene, with the exception of *L. shakespearei.* We categorized the *lpg1681*-containing loci into those with similar synteny, based on the orthology group annotations in Burstein et al. 2016. We colored and provided names for the neighboring genes in Supplemental figure 6 if they had a homolog in the *L. pneumophila* Philadelphia-1 genome that was not annotated as a hypothetical gene. To assess the conservation of the *lpg1680-lpg1682* among *L. pneumophila* strains, we used blastn in the NCBI nr and wgs databases with the full *lpg1680-lpg1682* genomic DNA sequence as a query. We found that the full region was conserved with few mutations across 501 currently sequenced *L. pneumophila* strains.

## Acknowledgements

We thank Howard Shuman for providing *Legionella* strains as well as the pLAW344 and pMMB207c plasmids, and Tamara O’Connor for the pTO100 plasmid carrying the transposon. Thank you to Michelle Swanson for providing the stringent response pathway mutant and complementation strains. We thank Ben Ross, Julian Simon, Rasi Subramaniam, Howard Shuman, Pete Greenberg, Jim Imlay, and the three reviewers for useful discussion and suggestions. We also thank Michelle Hays, Kevin Forsberg, Alistair Russell, Janet Young, and Howard Shuman for providing feedback on the manuscript.

---

**Supplemental Figure 1.** Separation of surfactant and antimicrobial phenotypes. A) Lm inhibition sometimes co-occurs with contact with WT Lp’s secreted surfactant (left), but sometimes is observable outside of the surfactant front (right). The leading edge of the surfactant front (arrowheads) is faintly visible when light is reflected off adjacent regions of the plate (“light”). Each image is a composite of two fields of view, with inverted colors to enable visualization of the surfactant front. B) Lack of surfactant production from Lp with a deletion of bbcB, evident by a defect in spreading on BCYE plates. C) The bbcB mutant can inhibit the growth of neighboring Lm, despite lacking surfactant secretion.

**Supplemental figure 2.** Genetic validation linking inhibition-defective mutants to the HGA-melanin pathway. A) Inter-bacterial inhibition does not correlate with surface spreading. For example, two mutants with transposon insertions in *lpg1874* (general secretion system protein L) and *lpg1875* (general secretion system protein M) share “small zone” inhibition phenotypes, yet show opposite spreading phenotypes on BCYE. B) Overexpression of either *hisC2* or *hpd* from a plasmid was sufficient to complement the “no zone” phenotype in all recovered mutants. Unlike mutations to *hisC2*, deletion of the *hmgA* gene does not disrupt *Lm* inhibition, showing that the intracellular recycling of HGA is not required for inhibitor production. Colors in images from B and C were inverted to facilitate visualization of the zone of inhibition. D) Pigmentation of AYE media following 48 hours growth of various *Legionella* species. While multiple species produce some pigment, many are less pigmented than *L. pneumophila*. The *L. micdadei* susceptible strain does not secrete detectable pigment. E) After 48 hours growth in AYE, none of the “no zone” mutants (represented here by *hisC2*::Tn) produce pigment. A subset of “small zone” mutants also have pigmentation defects (e.g. *lpg2872*::Tn and *lpg1875*::Tn), further implicating the HGA-melanin pathway in inhibition.

**Supplemental Figure 3.** Impacts of chemical compounds on HGA-mediated inhibition of *Legionella*. A) HGA inhibits *Lm* growth but HGA-related compounds do not. Chemical names and structures are shown above the corresponding plates. On each plate, different concentrations of each compound were spotted onto a lawn of *Lm* in 10µL droplets, arranged as indicated at the right. B) Addition of catalase does not rescue *Lm* susceptibility to 250 µM HGA. Control experiment showing catalase is active, based on its ability to protect *Lp* from H_2_O_2_. Quantification of HGA’s ability to oxidize cysteine. Data points are in purple, with a linear line of best fit. A 1:1 line is shown in gray, for reference. E) The potency of HGA is decreased when pre-incubated with reducing agents. 100mM HGA was mixed with dithiothreitol (DTT), oxidized glutathione (Ox. Glut), or reduced glutathione (R. Glut) for 15 minutes prior to spotting 10µL onto a lawn of *Lm* and allowing to grow for 3 days. Key below each image indicates where each solution was added.

**Supplemental Figure 4.** HGA-induced growth delays and quantification of HGA production. A-B) Growth curves of *Lp* strains upon exposure to HGA. For each experiment, we provide a matched wild type (KS79) control for comparison. A) Clinical isolate Philadelphia-1 and lab strain KS79 exhibited nearly identical growth curves in the presence of HGA. B) The Δ*hmgA* deletion strain responded to HGA similarly to wild type KS79, showing that HmgA-C do not play a significant role in HGA susceptibility. C) Viable CFU counts of KS79 *Lp* exposed to HGA. Even at high HGA concentrations, in rich media HGA is bacteriostatic at early time points, followed by population recovery. D-F) Using standard curves of synthetic HGA to estimate the amount of HGA secreted by *Lp*. D) OD400 was used to track the oxidation of synthetic HGA in AYE media. The 24 hour timepoint (arrow) was used for the standard curve and all experiments estimating HGA concentrations, as the HGA had completed oxidation by this point. E) Standard curve showing OD400 of oxidized synthetic HGA used to estimate the amount of HGA secreted into *Lp* conditioned media after 48 hours growth. The equation for the linear regression is shown along with a table with the OD400 readings and estimated HGA concentrations for each sample. F) Similar standard curve used for time course experiment of HGA secretion in wild type *Lp* in Figure 6. Standard curves were generated for each experiment independently, in parallel with experimental samples.

**Supplemental Figure 5.** Growth curves of lpg1681 Lp strains upon exposure to HGA. For each experiment we provide a matched wild type (KS79) control for comparison. The spontaneous lpg1681 mutant and the deletion mutant both had less severe growth delays from HGA than wild type. Conversely, the lpg1681 overexpression strain was sensitized to HGA, with longer growth delays than wild type. Dashed gray line shows time for the 250µM HGA condition to reach an OD of 0.1, to facilitate comparisons.

**Supplemental Figure 6.** Evolution of genes in the *lpg1681* locus among A) *Legionella* species and B) the fish pathogen *Piscirickettsia*. In the species that carry hypothetical gene *lpg1681*, it always resides upstream of the thiol:disulfide interchange gene *dsbD2*, despite extensive turnover of neighboring genes. Species separated by a “/” have similar syntenic loci. Annotated genes are colored, with redox-related genes in shades of blue. Hypothetical genes are white. Gray shading indicates gene homology among the species.

**Supplemental Figure 7.** Density-dependent HGA resistance acts independently of bulk cellular material, the *lqsR* quorum sensing pathway, and the stringent response pathway. A) The presence of dense heat-killed bacteria is not sufficient to protect dilute, live *Lp* cells from HGA inhibition. B) The susceptibility of wild type *Lp* (KS79) is nearly identical to that of *Lp* with a deletion of *lqsR*, the proposed quorum sensing response regulator in *L. pneumophila*. C) Schematic illustrating part of the stringent response pathway in *L. pneumophila*. In low nutrient conditions, RelA and SpoT generate the alarmone ppGpp, which activates a variety of downstream stress responses via the LetA/LetS two component system and the RpoS sigma factor. D) *L. pneumophila* stringent response mutants show similar, density-dependent susceptibility to HGA as wild type. Complementation of *relAspoT, rpoS*, and *letA* mutants by plasmid expression (with pSpoT, pRpoS, and pLetA respectively) similar had little effect on HGA inhibition.

**Supplemental table 1:** Transposon mutants of *L. pneumophila* with defects in *L. micdadei* inhibition. *The transposon insertions were mapped to a window that overlapped with both genes noted.

**Supplemental table 2:** Strains and plasmids

**Supplemental table 3:** Primers used for cloning

