## Supplemental Figures and Tables for "Density-dependent resistance protects *Legionella pneumophila* from its own antimicrobial metabolite, HGA"

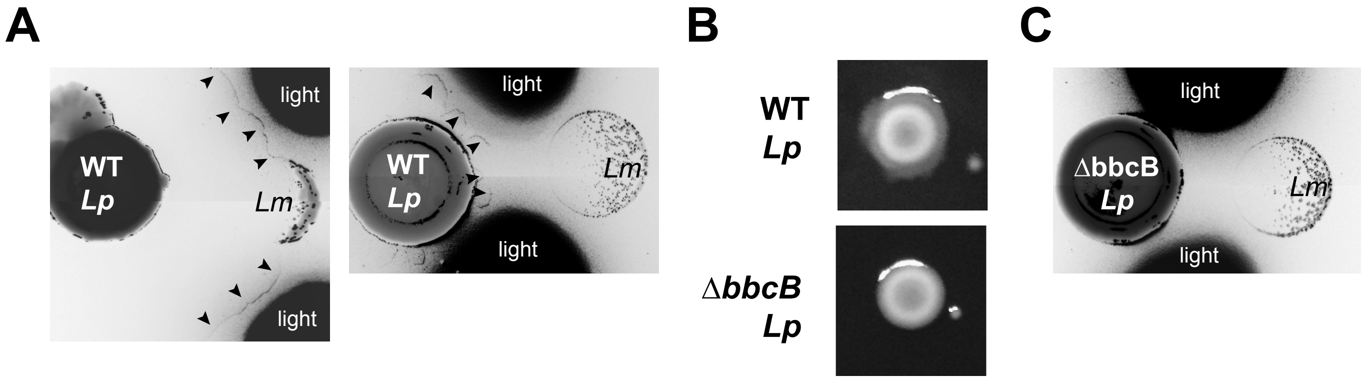


**Supplemental Figure 1.** Separation of surfactant and antimicrobial phenotypes.
**A)** *Lm* inhibition sometimes co-occurs with contact with WT *Lp*'s secreted surfactant (left), but sometimes is observable outside of the surfactant front (right). The leading edge of the surfactant front (arrowheads) is faintly visible when light is reflected off adjacent regions of the plate (”light”). Each image is a composite of two fields of view, with inverted colors to enable visualization of the surfactant front. **B)** Lack of surfactant production from *Lp* with a deletion of *bbcB*, evident by a defect in spreading on BCYE plates. **C)** The *bbcB* mutant can inhibit the growth of neighboring *Lm*, despite lacking surfactant secretion.

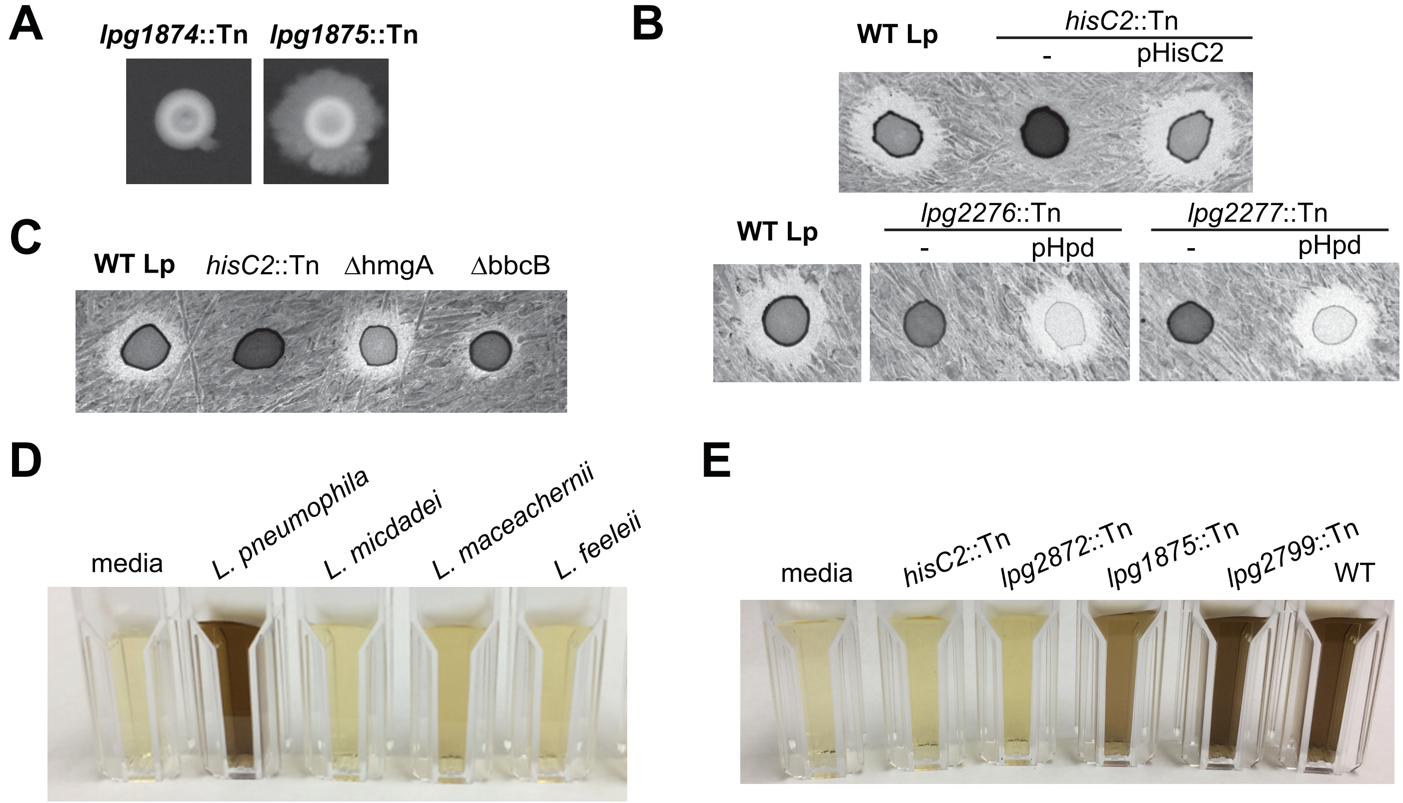


**Supplemental figure 2.** Genetic validation linking inhibition-defective mutants to the HGA-melanin pathway. **A)** Inter-bacterial inhibition does not correlate with surface spreading. For example, two mutants with transposon insertions in *lpg1874* (general secretion system protein L) and *lpg1875* (general secretion system protein M) share "small zone" inhibition phenotypes, yet show opposite spreading phenotypes on BCYE. **B)** Overexpression of either *hisC2* or *hpd* from a plasmid was sufficient to complement the "no zone" phenotype in recovered mutants. **C)** Unlike mutations to *hisC2*, deletion of the *hmgA* gene does not disrupt *Lm* inhibition, showing that the intracellular recycling of HGA is not required for inhibitor production. Colors in images from B and C were inverted to facilitate visualization of the zone of inhibition. **D)** Pigmentation of AYE media following 48 hours growth of various *Legionella* species. While multiple strains produce some pigment, many are less pigmented than *L. pneumophila*. The *L. micdadei* susceptible strain does not secrete detectable pigment. **E)** After 48 hours growth in AYE, none of the "no zone" mutants (represented here by *hisC*2::Tn) produce pigment. A subset of "small zone" mutants also have pigmentation defects (e.g. *lpg2872*::Tn and *lpg1875*::Tn), further implicating the HGA-melanin pathway in inhibition.

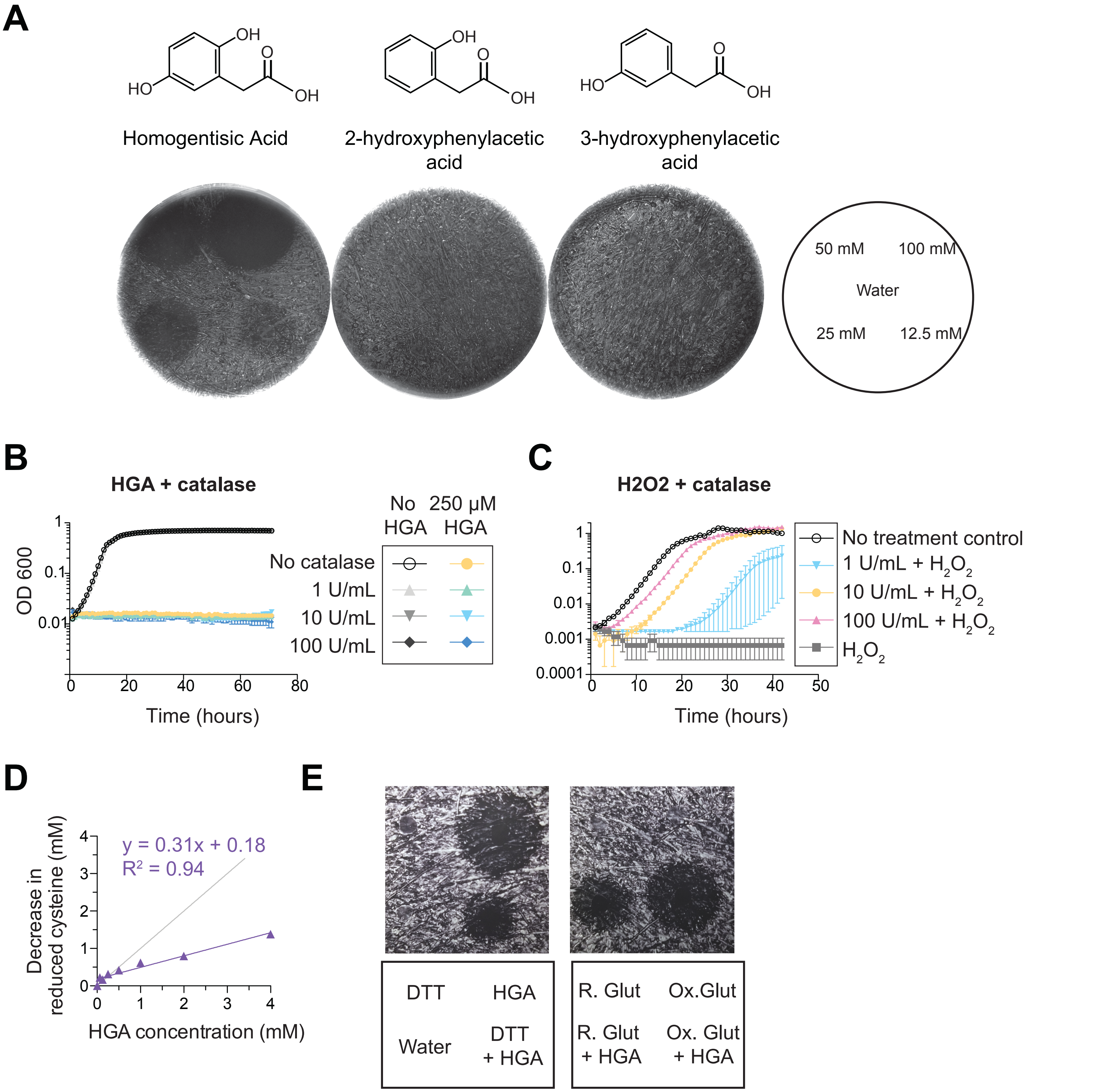


**Supplemental Figure 3**. Impacts of chemical compounds on HGA-mediated inhibition of *Legionella*.
**A**) HGA inhibits *Lm* growth but HGA-related compounds do not. Chemical names and structures are shown above the corresponding plates. On each plate, different concentrations of each compound were spotted onto a lawn of *Lm* in 10 μL droplets, arranged as indicated at the right. **B)** Addition of catalase does not rescue *Lm* susceptibility to 250 μM HGA. **C)** Control experiment showing catalase is active, based on its ability to protect *Lp* from $H_2O_2$. **D)** Quantification of HGA's ability to oxidize cysteine. Data points are in purple, with a linear line of best fit. A 1:1 line is shown in gray, for reference. **E)** The potency of HGA is decreased when pre-incubated wtih reducing agents. 100 mM HGA was mixed with dithiothreitol (DTT), oxidized glutathione (Ox. Glut) or reduced glutathione (R. Glut) for 15 minutes prior to spotting 10 μL onto a lawn of *Lm* and allowing to grow for 3 days. Key below each image indicates where each solution was added.

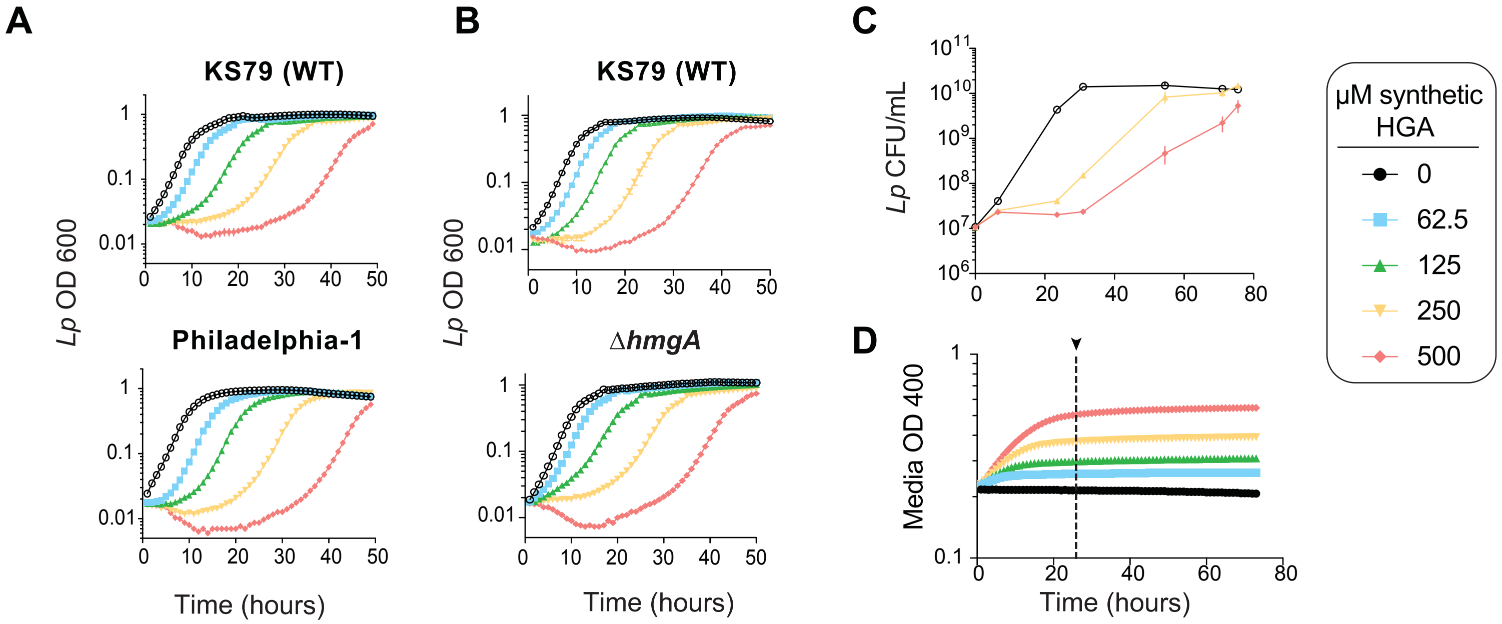


E

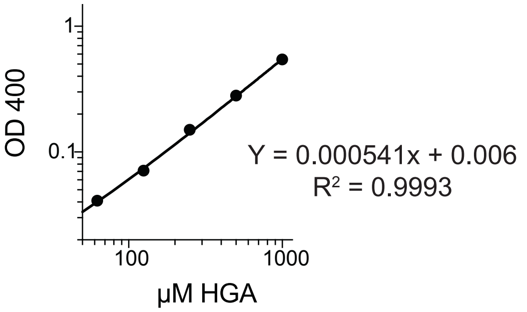


| *Lp* strain | Oxidized media OD 400 | Estimated [HGA] (μM) |
|---|---|---|
| WT | 0.91 | 1.7 mM |
| Δ*hmgA* | 1.39 | 2.6 mM |
| *hisC2*::Tn | 0 | 0 mM |

F

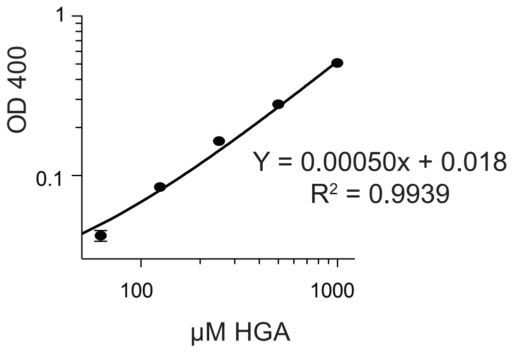


| Time (hours) | Oxidized media OD 400 | Estimated [HGA] (μM) |
|---|---|---|
| 0 | 0 | <62.5 |
| 8 | -0.03 | <62.5 |
| 24 | 0 | <62.5 |
| 29 | 0.11 | 183 |
| 32 | 0.24 | 448 |
| 48 | 0.34 | 648 |

**Supplemental Figure 4**. HGA-induced growth delays and quantification of HGA production. **A-B)** Growth curves of *Lp* strains upon exposure to HGA. For each experiment, we provide a matched wild type (KS79) control for comparison. **A)** Clinical isolate Philadelphia-1 and lab strain KS79 exhibited nearly identical growth curves in the presence of HGA. **B)** The Δ*hmgA* deletion strain responded to HGA similarly to wild type KS79, showing that HmgA-C do not play a significant role in HGA susceptibility. **C)** Viable CFU counts of KS79 *Lp* exposed to HGA. Even at high HGA concentrations, in rich media HGA is bacteriostatic at early time points, followed by population recovery. D-F) Using standard curves of synthetic HGA to estimate the amount of HGA secreted by *Lp*. **D)** OD400 was used to track the oxidation of synthetic HGA in AYE media. The 24 hour timepoint (arrow) was used for the standard curve and all experiments estimating HGA concentrations, as the HGA had completed oxidation by this point. **E)** Standard curve showing OD400 of oxidized synthetic HGA used to estimate the amount of HGA secreted into *Lp* conditioned media after 48 hours growth. The equation for linear regression is shown along with a table with the OD400 readings and estimated HGA concentrations for each sample. **F)** Similar standard curve used for time course experiment of HGA secretion in wild type *Lp* in Figure 6. Standard curves were generated for each experiment independently, in parallel with experimental samples.

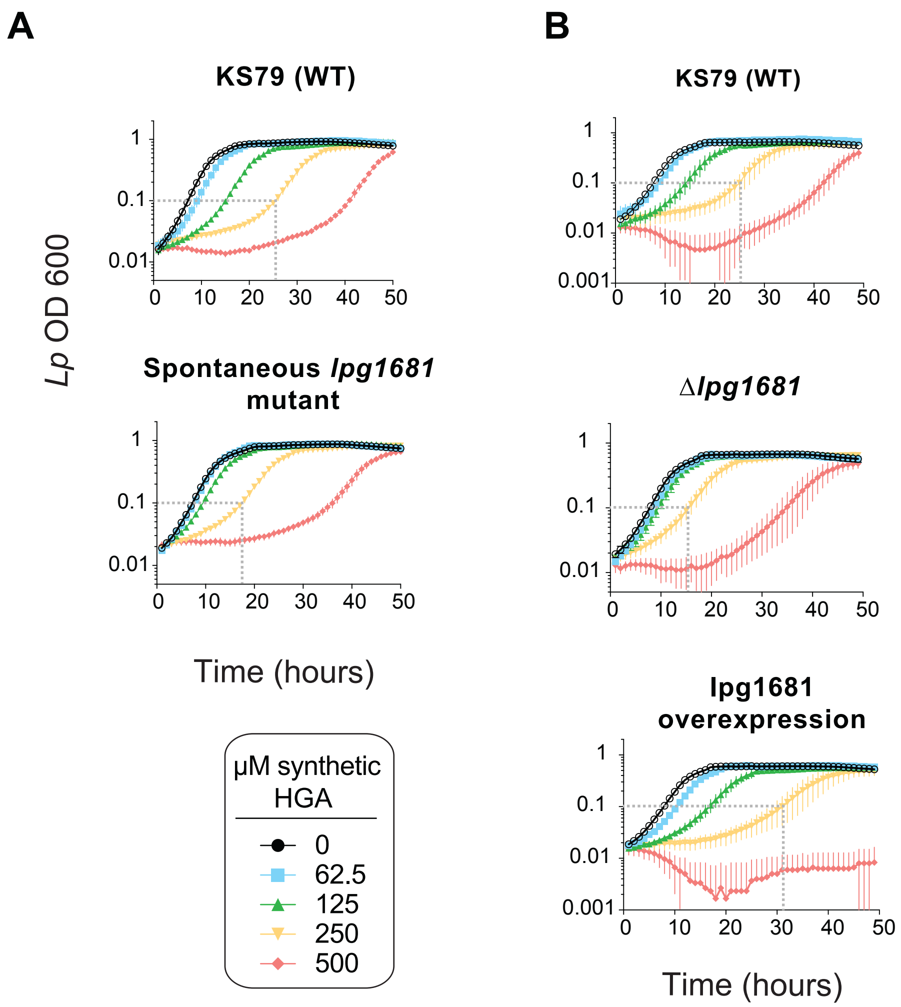


**Supplemental Figure 5.** Growth curves of *lpg1681 Lp* strains upon exposure to HGA. For each experiment, we provide a matched wild type (KS79) control for comparison. The spontaneous *lpg1681* mutant and the deletion mutant both had less severe growth delays from HGA than wild type. Conversely, the *lpg1681* overexpression strain was sensitized to HGA, with longer growth delays than wild type. Dashed gray line shows the time for the 250µM HGA condition to reach an OD of 0.1, to facilitate comparisons.

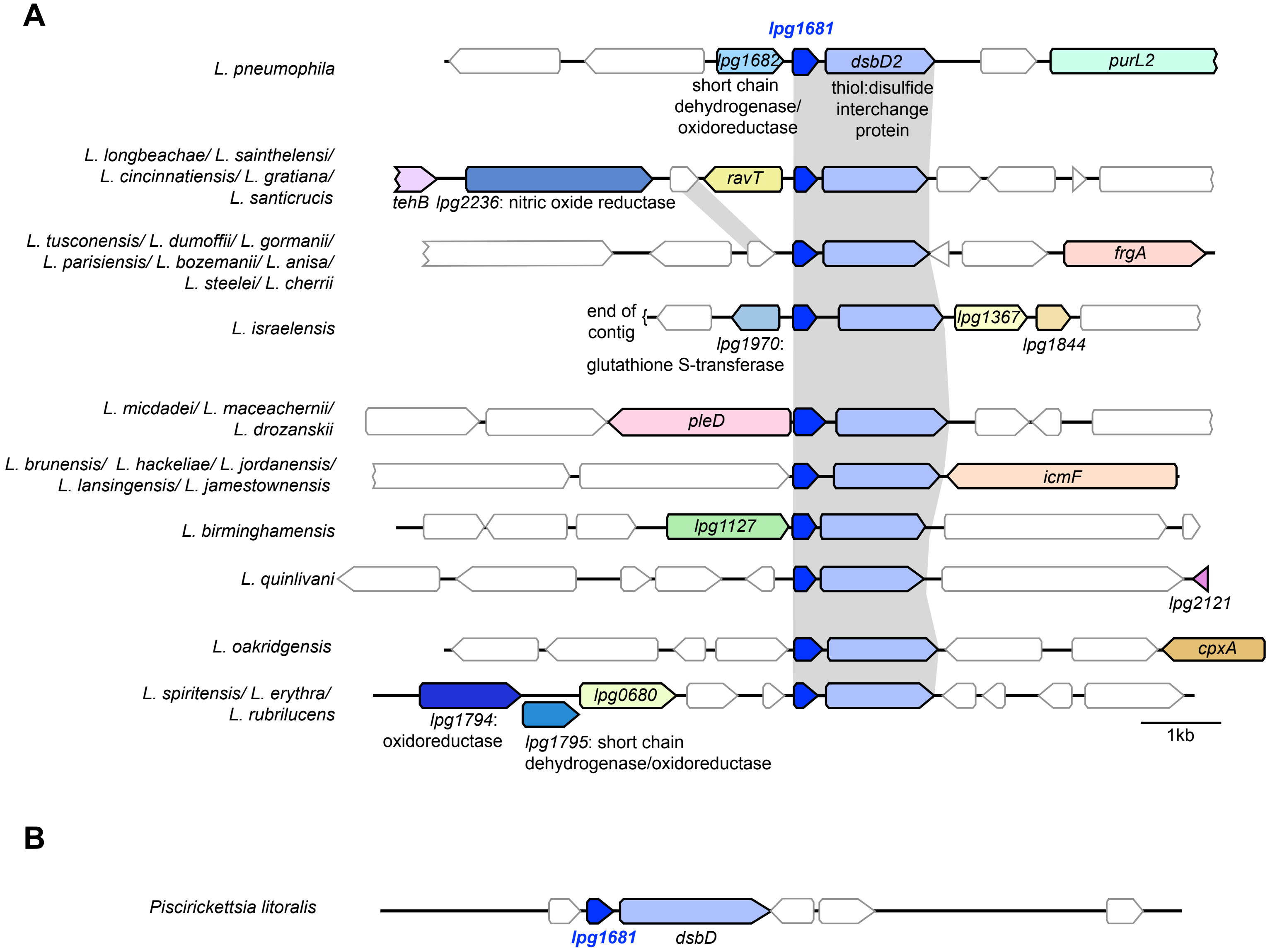


**Supplemental Figure 6.** Evolution of genes in the *lpg1681* locus among **A)** *Legionella* species and **B)** the fish pathogen *Piscirickettsia*. In the species that carry hypothetical gene *lpg1681*, it always resides upstream of the thiol:disulfide interchange gene *dsbD2*, despite extensive turnover of neighboring genes. Species separated by a "/" have similar syntenic loci. Annotated genes are colored, with redox-related genes in shades of blue. Hypothetical genes are white. Gray shading indicates gene homology among the species.

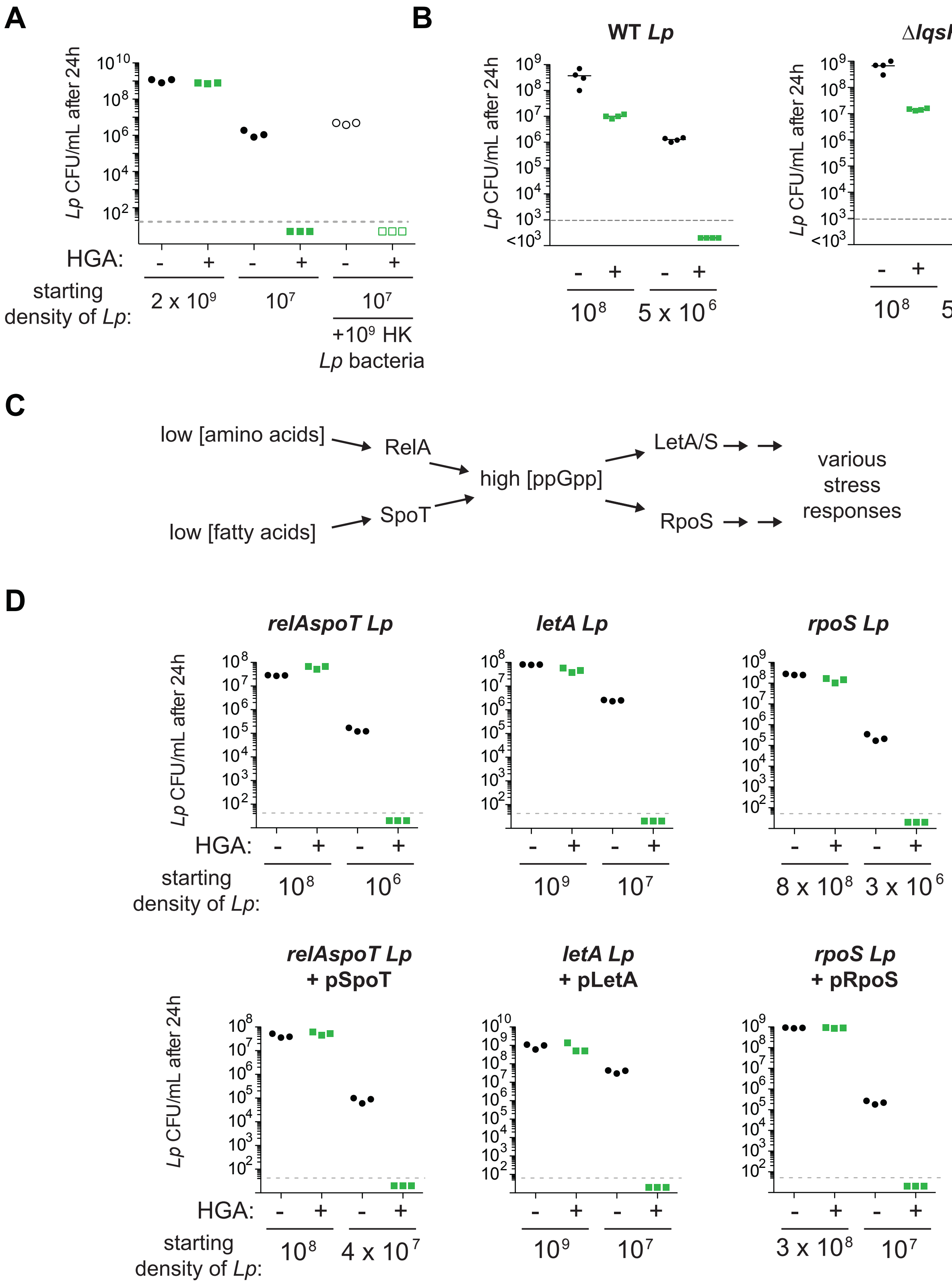


**Supplemental Figure 7.** Density-dependent HGA resistance acts independently of bulk cellular material, the *lqsR* quorum sensing pathway, and the stringent response pathway. **A)** The presence of dense heat-killed bacteria is not sufficient to protect dilute, live *Lp* cells from HGA inhibition. **B**) The susceptibility of wild type *Lp* (KS79) is nearly identical to that of *Lp* with a deletion of *lqsR*, the proposed quorum sensing response regulator in *L. pneumophila*. **C**) Schematic illustrating part of the stringent response pathway in *L. pneumophila*. In low nutrient conditions, RelA and SpoT generate the alarmone ppGpp, which activates a variety of downstream stress responses via the LetA/LetS two component system and the RpoS sigma factor. **D**) *L. pneumophila* stringent response mutants show similar, density-dependent susceptibility to HGA as wild type. Complementation of *relAspoT, rpoS*, and *letA* mutants by plasmid expression (with pSpoT, pRpoS, and pLetA respectively) similarly had little effect on HGA inhibition.

Table 1: Genes Mutated in HGA-Selected *L. pneumophila*

| **Mutated locus** | **Function or Product** | **# Spontaneous Mutants w/ this mutation** | **Mutations Recovered** |
|---|---|---|---|
| *lpg1681* | Hypothetical Protein | 4 | R49K, R49S, R49G, T50K |
| *lpg0288* | YjeK, 2,3-beta-lysine aminomutase | 11 | W8S, Q9*, L26G, K39*, A50P, R101L, D196G, H215R, Q243K, I250F, Q294* |
| *lpg0607* | PoxA/YjeA/GenX, Elongation factor P beta-lysine transferrase | 5 | W100*, Q184L, A218V, Q258*, 1 bp deletion in S156 |
| *lpg0325* | RpS7 ribosomal 30S protein | 1 | G100D |
| *lpg0336* | RplP 50s ribosomal protein | 1 | G88R |
| *lpg0287* | Elongation factor P | 1 | Stop>Q (70 AA Extension) |
| *lpg0349* | SecY | 3 | N118K, Q132R, R369L |
| *lpg2001* | SecD | 2 | V238F, A277G |
| *lpg1504* | AceE pyruvate dehydrogenase | 1 | P272C |

* =introduction of a stop codon

Supplemental table 1: Transposon mutants of *L. pneumophila* with defects in *L. micdadei* inhibition

| Protein Group and Locus of Insertion | Function or Product Name | Polar Effect Possible? | Upstream of Transport Gene? | Hits in the Same Operon | Pigment Production | Inhibition Phenotype |
|---|---|---|---|---|---|---|
| *lpg1998* | Histidinol-Phosphate Aminotransferase (hisC2) | Yes | No | 2 | None | No Zone |
| *lpg2276* | Glu/Leu/Phe/Val Dehydrogenase | Yes | No | 3 | None | No Zone |
| *lpg2277* | O-methyltransferase, SAM-Dependent | Yes | No | 3 | None | No Zone |
| *lpg0501* | Phosphoglyceromutase (pgm) | Yes | No | 1 | Yes | Small Zone |
| *lpg0675* | Hypothetical gene | No | No | 1 | Yes | Small Zone |
| *lpg0710* | Sepiapterin reductase (yueD) | Yes (hutI) | Yes | 2 | None | Small Zone |
| *lpg0711* | Hydrolase of 4-imidazolone-5-propionate (hutI) | Yes | Yes | 2 | Reduced | Small Zone |
| *lpg1020* | Chemiosmotic Efflux System B Protein A | No | Yes | 1 | Yes | Small Zone |
| *lpg1291* | 2 component sensor kinase | Yes | No | 1 | Yes | Small Zone |
| *lpg1439* | Mg2+ and Co2+ Transporter (corC) | Yes | Yes | 1 | Yes | Small Zone |
| *lpg1867* | Phage Integrase | Yes | Yes | 1 | Yes | Small Zone |
| *lpg1874* | General secretion pathway protein L | Yes (yghD) | Yes | 2 | Yes | Small Zone |
| *lpg1875* | General secretion pathway protein M-type (yghD) | No | Yes | 2 | Yes | Small Zone |
| *lpg2238/ lpg2239** | Hypothetical gene | Yes | Yes | 1 | Yes | Small Zone |
| *lpg2248* | Hypothetical gene | Yes | No | 2 | Yes | Small Zone |
| *lpg2251* | Hypothetical gene | Yes | No | 2 | Yes | Small Zone |
| *lpg2269* | Endonuclease | Yes | No | 1 | Yes | Small Zone |

| | | | | | | |
|---|---|---|---|---|---|---|
| *lpg2439* | NADPH-dependent FMN reductase domain-containing protein | No | No | 1 | Yes | Small Zone |
| *lpg2478/ lpg2477** | glycotransferase or high affinity nickel transporter | Yes | Yes | 1 | Yes | Small Zone |
| *lpg2679* | D-isomer specific 2-hydroxyacid dehydrogenase | Yes | No | 1 | Yes | Small Zone |
| *lpg2799* | O-acetyltransferase | No | No | 1 | Yes | Small Zone |
| *lpg2872* | dinucleoside phosphate hydrolase | Yes | No | 1 | Reduced | Small Zone |

* The transposon insertions were mapped to a window that overlapped with both genes noted

Supplemental table 2: Strains, plasmids, and primers

| **Strains** | **Relevant Genotype** | **References** |
|---|---|---|
| KS79 | Wild-type *Legionella pneumophila (Lp)* | de Felipe *et al*, 2008 |
| Tatlock | Wild-type *Legionella micdadei (Lm)* | Herbert *et al.*, 1980 |
| LG36 | KS79 Δ*bbcB* | This work |
| LG51 | *hisC2*::Tn insertion in KS79 | This work |
| LG54 | *lpg2276*::Tn in KS79 | This work |
| LG79 | *lpg2277*::Tn in KS79 | This work |
| LG38 | KS79 Δ*hmgA* | This work |
| LG102 | KS79 Δ*lpg1681* | This work |
| LG107 | KS79 Δ*lqsR* | This work |
| MB444 | Lp02 *rpoS* pRpoS-pKB5 | Bachman and Swanson, 2001 |
| MB445 | Lp02 *rpoS* pKB5 | Bachman and Swanson, 2001 |
| MB434 | Lp02 *letA* pMMB | Hammer *et al.*, 2002 |
| MB435 | Lp02 *letA* pLetA-pMMB | Hammer *et al.*, 2002 |
| MB697 | Lp02 *relA*:gent/*spoT*:kan | Dalebroux *et al.*, 2008 |
| MB688 | Lp02 *relA*:gent/*spoT*:kan/pspoT | Dalebroux *et al.*, 2008 |

| **Plasmids** | **Description** | |
|---|---|---|
| pMMB207C | RSF1010 derivative, IncQ lacI$^q$ Cm$^r$ P*tac* *ori*T Δ*mobA* | Chen *et al.*, 2004 |
| pLAW344 | SacB MCS *oriT*(RK2) Cm$^r$ *oriR*(Co1E1) Ap$^r$ *loxP* | Wiater *et al.*, 1994 |
| pTO100 | *Mariner himar*-1 transposon R6K Km$^r$ *sacB* Ap$^r$ C9 transposase | O'Connor *et al.,* 2011 |
| pTL35 | pLAW344:ΔbbcB | This work |
| pTL36 | pLAW344:ΔhmgA | This work |
| pTL43 | pMMB207c:hisC2 | This work |
| pTL46 | pMMB207c:Hpd | This work |
| pTL52 | pLAW344:Δ1681 | This work |
| pTL53 | pLAW344:ΔLqsR | This work |

| | | |
|---|---|---|
| pTL54 | pMMB207c:lpg1681 | This work |

| **Gene Deletion Primers** | Sequence (5’ -> 3’) |
|---|---|
| *bbcB* P1 | TCCACTAGTTCTAGAGCGGCCCAGACAAGATAAGGCCACT |
| *bbcB* P2 | AAAAACAGCATGGTTTTATTCCTCAATATAAAACACCTTAATTGCTATATGATGCTTT |
| *bbcB* P3 | AAAGCATCATATAGCAATTAAGGTGTTTTATATTGAGGAATAAAACCATGCTGTTTTT |
| *bbcB* P4 | GGAAGATACTTAACAGGGAAGTGAGAGTTCAGAAATGGAAGAAGGCAG |
| *hmgA* P1 | TCCACTAGTTCTAGAGCGGCCAAGATGATGAAGAGCCAATTAAGGA |
| *hmgA* P2 | CATCAAAAATCTTGAAGGAGATTGAACTATTGACTGCTTTTTAAGATGTAAAATTGAC |
| *hmgA* P3 | GTCAATTTTACATCTTAAAAAGCAGTCAATAGTTCAATCTCCTTCAAGATTTTTGATG |
| *hmgA* P4 | GGAAGATACTTAACAGGGAAGTGAGACTTAATTCCGTATCGTTTGATGTGCG |
| *1681* P1 | TCCACTAGTTCTAGAGCGGCCCCAATCCCTGAGCCAAATAACC |
| *1681* P2 | CTGTACAGACCTCTCTTAAAAAGATACTATAGAGTAAATTTGTCAACTTTCTCAGTAGG |
| *1681* P3 | CCTACTGAGAAAGTTGACAAATTTACTCTATAGTATCTTTTTAAGAGAGGTCTGTACAG |
| *1681* P4 | GGAAGATACTTAACAGGGAAGTGAGAGGCAATTTCTCATCACCTTCAAG |
| *LqsR* P1 | TCCACTAGTTCTAGAGCGGCCAACATCAGTCAACATCCGCAA |
| *LqsR* P2 | ACTCCAAAAAGCCCGGTGAAAACTGTTCAAATGAGCCGCC |
| *LqsR* P3 | GGCGGCTCATTTGAACAGTTTTCACCGGGCTTTTTGGAGT |
| *LqsR* P4 | GGAAGATACTTAACAGGGAAGTGAGAGCACAGCGTGGGGAGATTAA |
| **Complementation Primers** | |
| *hpd* Fwd | CAATTTCACACAGGAAACAGAATCCCCGGCAAGCAATACA |
| *hpd* Rev | TGCCTGCAGGTCGACTCTAGCTAGCTTAATTCTTTTAAAGTACCACGTCGAAC |
| *hisC2* Fwd | GCTCGGTACCCGGGGATCCTATAAATTCATTCGTGACAGCTGGC |
| *hisC2* Rev | TGCCTGCAGGTCGACTCTAGCGCCAATTCCCTCGCAAG |
| *lpg1681* Fwd | CAATTTCACACAGGAAACAGGACCTCTCTTAAAAAGATACTATAGTTATTATG |
| *lpg1681* Rev | TGCCTGCAGGTCGACTCTAGGCCTACTGAGAAAGTTGACAA |
| **Arbitrary PCR Primers** | |
| **pTO100_F** | CTTCTATCGCCTTCTTGACGAGTT |
| **pTO100_R** | GTGCCACCTAAATTGTAAGCGTTA |

| | |
|---|---|
| **pTO100_Rd2** | TCTAGAGACCGGGGACTTATCA |
| **arb1** | GGCCACGCGTGCACTAGTACNNNNNNNNNNGATAT |
| **arb2** | GGCCACGCGTCGACTAGTAC |
